# A tryptophan metabolite modulates the host response to bacterial infection via kainate receptors

**DOI:** 10.1101/2023.08.16.553532

**Authors:** Margarita Parada-Kusz, Anne E. Clatworthy, Emily R. Goering, Stephanie M. Blackwood, Elizabeth J. Salm, Catherine Choi, Senya Combs, Jenny S. W. Lee, Carlos Rodriguez-Osorio, Susumu Tomita, Deborah T. Hung

## Abstract

Bacterial infection involves a complex interaction between the pathogen and host where the outcome of infection is not solely determined by pathogen eradication. To identify small molecules that promote host survival by altering the host-pathogen dynamic, we conducted an *in vivo* chemical screen using zebrafish embryos and found that treatment with 3-hydroxy-kynurenine protects from lethal gram-negative bacterial infection. 3-hydroxy-kynurenine, a metabolite produced through host tryptophan metabolism, has no direct antibacterial activity but enhances host survival by restricting bacterial expansion in macrophages by targeting kainate-sensitive glutamate receptors. These findings reveal new mechanisms by which tryptophan metabolism and kainate-sensitive glutamate receptors function and interact to modulate immunity, with significant implications for the coordination between the immune and nervous systems in pathological conditions.

## INTRODUCTION

Since the discovery of antibiotics almost a century ago, the predominant therapeutic strategy in addressing bacterial infection is to identify molecules that kill or restrict the growth of bacteria *in vitro* and then translate these antibiotics for *in vivo* use. Yet this approach ignores the complex dynamic between pathogen and host, and the numerous mechanisms involved in their interplay that can contribute to determining infection outcome (*1, 2*). To elucidate critical mechanisms in this interaction that contribute to control of infection, we sought to identify small molecules that restore host health, not by killing bacteria directly, similar to antibiotics, but by altering the host-pathogen dynamic to favor the host. In order to identify molecules that act in this way, we performed whole-organism chemical screening using zebrafish embryos as a model vertebrate host, as their immune system is similar to mammals (*3*) and they are amenable to both classical and chemical genetics. We identified that exogenous addition of the tryptophan metabolite 3-hydroxy-kynurenine (3-HK) protects against lethal, systemic gram-negative bacterial infections by *Pseudomonas aeruginosa* and *Salmonella enterica* serovar Typhimurium (*S.* Typhimurium).

3-HK is endogenously produced in all eukaryotes as an intermediate metabolite in the kynurenine pathway in which tryptophan is oxidized to NAD+ (*4*), with several metabolites in this pathway implicated in neurodegenerative diseases, inflammatory disorders and cancer (*4–7*). We found that in addition to exogenous supplementation of 3-HK protecting against systemic infection with gram-negative pathogens, inhibition of its endogenous production results in sensitization to infection. 3-HK restricts bacterial expansion in macrophages in the context of the whole organism and promotes survival by acting at ionotropic glutamate receptors (iGluRs) of the kainate-sensitive (KAR) subclass, which are ligand-gated ion channels primarily studied for their role in modulating synaptic transmission in vertebrates. Importantly, while other kynurenine pathway metabolites are known to interact with other iGluR subclasses to modulate neuronal excitability (*5, 7, 8*), this work expands the relationship between the kynurenine pathway and iGluRs to reveal a novel mechanism by which the kynurenine pathway acts in defense against infection, a role for KARs in immune defense, and a novel intersection between the two in host immunity.

## RESULTS

### 3-HK promotes survival following infection but does not act like a typical antibiotic

We screened a library of 932 known bioactive compounds for small molecules that promote zebrafish embryo survival following lethal *P. aeruginosa* challenge (*9*) and found that the small molecule 3-HK rescued embryos infected with *P. aeruginosa* when added to the aqueous media surrounding infected fish (Fig. 1A). 3-HK was even more potent against infection by the gram-negative pathogen *S.* Typhimurium (Fig. 1B) but had little effect on lethal *Staphylococcus aureus* or *Mycobacterium marinum* challenge (fig. S1). 3-HK significantly repressed bacterial burden in infected fish over time, as can be achieved with the antibiotic ciprofloxacin (Fig. 1C). However, 3-HK does not work like a traditional antibiotic as 3-HK had no effect on *P. aeruginosa* or *S.* Typhimurium axenic growth *in vitro* in either rich (LB) or minimal (M9) media (Fig. 1D and fig S2A-C). Given the potent survival benefit that 3-HK had against *S.* Typhimurium infection, we focused on *S.* Typhimurium in all subsequent studies of 3-HK.

**Fig. 1.**
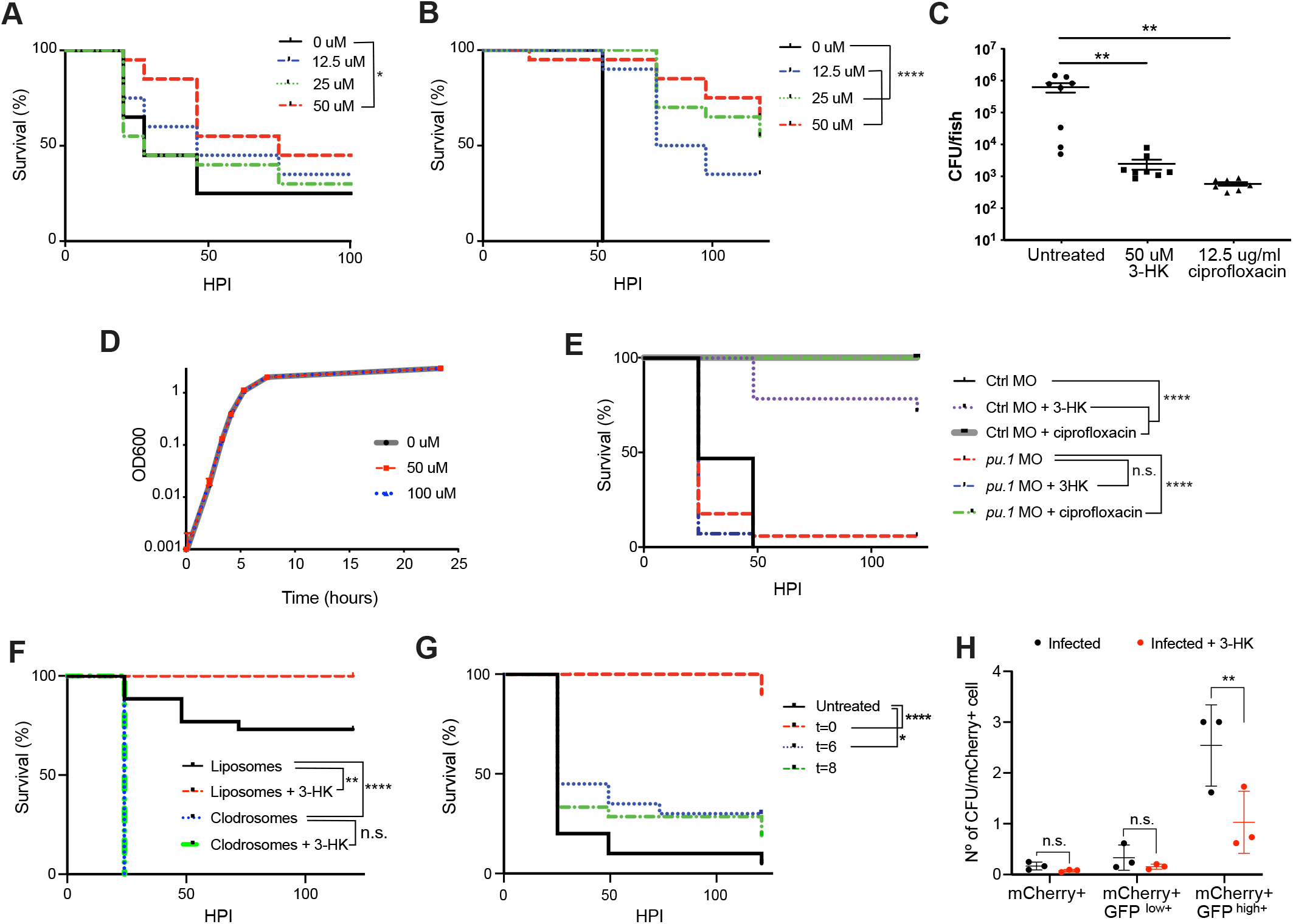
3-HK promotes survival following infection but does not act like a classical antibiotic. Embryos (48 hpf; N=20/condition) were infected intravenously with either *P. aeruginosa* (**A**; 9095+/-1034 CFU) or *S.* Typhimurium (**B**; 333+/-53 CFU) and treated with indicated concentrations of 3-HK, which significantly promoted survival ((**A**) *, P=0.0391 and (**B**) ****, P<0.0001, log-rank test). (**C**) Larvae (72 hpf) were infected with *S.* Typhimurium (336+/-86 CFU), treated with vehicle control, 3-HK or the antibiotic ciprofloxacin and bacterial burden was enumerated from individual fish 19 hpi (one-way ANOVA, P=0.0012; Dunnett’s multiple comparisons test; **, P=0.0023, 3-HK treated; **, P=0.0022, cipro treated); data are representative of >3 independent experiments. (**D**) 3-HK does not inhibit growth of *S.* Typhimurium cultured *in vitro* in LB media. (**E**) *pu.1* or Control morphants (72 hpf) were infected with *S.* Typhimurium (342+/-37 CFU), treated with vehicle control, 50uM 3-HK or 12.5ug/ml cipro, and monitored over time for survival. 3-HK’s survival activity requires the presence of immune cells, unlike cipro (N=14-17 larvae/ condition; ****, P< 0.0001, log-rank test). (**F**) Larvae (72 hpf) were injected with 2.5 ng of either control liposomes or liposomes containing clodronate (*i.e.,* clodrosomes), to deplete macrophage numbers, and then infected with *S.* Typhimurium (447+/-23 CFU) and either treated with vehicle control or 50uM 3-HK and monitored over time for survival. 3-HK’s survival activity requires the presence of macrophages (N=24-26 larvae/ condition; **, P=0.0067; ****, P< 0.0001; log-rank test). (**G**) Larvae (72 hpf) were infected with 612+/-58 CFU, treated with 50 uM 3-HK at indicated hours post infection and monitored over time for survival (N=17-21 embryos/ condition; ****, P<0.0001; *, P=0.0352; log rank test). (**H**) *mpeg1:mCherry* larvae were infected with S.Tm-GFP (662+/-13 CFU), treated or untreated with 3-HK 50 uM, and at 4 hpi mCherry^+^ cells were sorted based upon GFP intensity emission as indicated, and plated for CFU and normalized according to the total number of cells recovered per condition (two-way ANOVA, **, P=0.0028, Šidák multiple comparison’s test. N= 3 ind. exps.). Survival curves data are representative of at least 3 independent experiments.

The activity of 3-HK was dependent on the presence of macrophages, a known favored replicative niche for *S.* Typhimurium in mammals and zebrafish (*10–12*), further distinguishing its mechanism of action from traditional antibiotics. Using a morpholino to block the expression of the transcription factor Pu.1, which shifts blood cell development away from myelopoiesis and toward erythropoiesis (*13*), we decreased the number of immune cells available to combat infection, which at this developmental stage consists of macrophages and neutrophils (fig. S3A-B)(*3*). While *pu.1* morphant fish were exquisitely sensitive to infection as previously described (*9, 14*), the activity of 3-HK also was significantly attenuated, in contrast to the broad-spectrum antibiotic ciprofloxacin which still effectively rescued infected fish (Fig. 1E)). Because knock-down of *pu.1* depletes both macrophages and neutrophils, we more selectively eliminated macrophages using clodronate liposomes (79). Like genetic manipulation of *pu.*1, clodronate liposomes also abrogated the effect of 3-HK, but not ciprofloxacin, on survival (Fig. 1F and fig. S3C-E). These results demonstrated that macrophages are required for the 3-HK effect.

In addition to the requirement of macrophages for 3-HK’s protective effect, there was also a time dependency for its administration. We found that 3-HK had full efficacy if administered early, up to 4 hours post infection (hpi) (fig. S4), but gradually lost efficacy when administered at later time points; 3-HK treatment at 8 hours post infection showed no survival benefit (Fig. 1G and fig. S4). As the vast majority of *S.* Typhimurium reside within macrophages at these early time points (*12, 15*), we examined whether 3-HK restricted bacterial expansion within macrophages at 4 hpi. We quantified bacterial burden by plating for colony forming units (CFU) from bacteria-containing macrophages sorted from macrophage transgenic reporter larvae (*mpeg1:mCherry*) infected with *S.* Typhimurium constitutively expressing GFP (S.Tm-GFP). We compared CFUs/macrophage from fish treated with and without 3-HK. Flow cytometry of mCherry^+^ macrophages from infected fish revealed a negative subpopulation and two subpopulations with different intensities of GFP fluorescence emission (negative, low and high; fig S5). The two levels of GFP intensity in *S.* Typhimurium infected macrophages is consistent with previous observations describing one group of macrophages which have successfully degraded intracellular bacteria thus having faint fluorescent signal (reminiscent of GFP^low+^), while another group of macrophages has live, persisting and/or dividing intracellular bacteria (reminiscent of GFP^high+^) (fig S6) (*12*). While 3-HK treatment had no impact on the total numbers of recovered macrophages for each category (fig S5B-E), 3-HK treated animals contained on average ∼50% less recoverable CFUs/macrophage in the GFP^high+^ cell population than untreated animals (Fig. 1H). These results demonstrate that 3-HK treatment restricts bacterial expansion in macrophages *in vivo*. In contrast, in more reductionist models of infection using cultured macrophage cell lines (J774, RAW264, or U397) or primary cells (murine C57BL/6 bone-marrow derived macrophages), 3-HK had no effect on host cell survival or bacterial burden (Table S1), suggesting that it may exert its protective effect indirectly or in a way not easily modeled outside the whole organism.

Taken together, the absence of 3-HK *in vitro* antibacterial activity, its dependence on macrophages for *in vivo* activity, and its ability to restrict intracellular bacteria only within the context of the whole organism, indicate that 3-HK does not directly target the bacterium like a classical antibiotic. Rather, 3-HK disrupts the pathogen-host dynamic, possibly by targeting the host by a more systemic mechanism.

### Host endogenous production of 3-HK is beneficial against *S.* Typhimurium infection

3-HK is a metabolite produced from tryptophan by the kynurenine pathway in all eukaryotes (*4*). While a form of the kynurenine pathway is also present in some prokaryotic species (*4*), including *P. aeruginosa* (*16, 17*) but not *S.* Typhimurium, neither bacterial species synthesizes 3-HK itself (Fig. 2A, fig. S7A)(*16, 17*). Since the prokaryotic kynurenine pathway is not conserved in both *P. aeruginosa* and *S.* Typhimurium and we hypothesized that 3-HK may be mediating its effects via the host, we focused on the host kynurenine pathway (Fig. 2A). In mammals, tryptophan is catabolized to N-formyl-kynurenine by three enzymes: tryptophan 2,3-dioxygenase (TDO); indoleamine 2,3-dioxygenase 1 (IDO1), which is IFN-*γ-*inducible and most highly expressed in professional antigen presenting cells (*e.g.*, macrophages and dendritic cells) and mucosal tissues; and indoleamine 2,3-dioxygenase 2 (IDO2)(*4, 8*). N-formyl-kynurenine is then hydrolyzed to kynurenine, which is then catabolized to 3-HK by the enzyme kynurenine 3-monooxygenase (KMO)(*4, 8*). KMO can accept either L-kynurenine or D-kynurenine as a substrate, and preserves the stereochemistry of the substrate in the product (*18*), yielding either 3-hydroxy-L-kynurenine or 3-hydroxy-D-kynurenine, respectively (Fig. 2A). 3-HK can then be further metabolized to 3-hydroxy-anthranilic acid (3-HAA) and then quinolinic acid (QA), with metabolism of QA ultimately resulting in the *de novo* synthesis of NAD^+^.

**Fig. 2.**
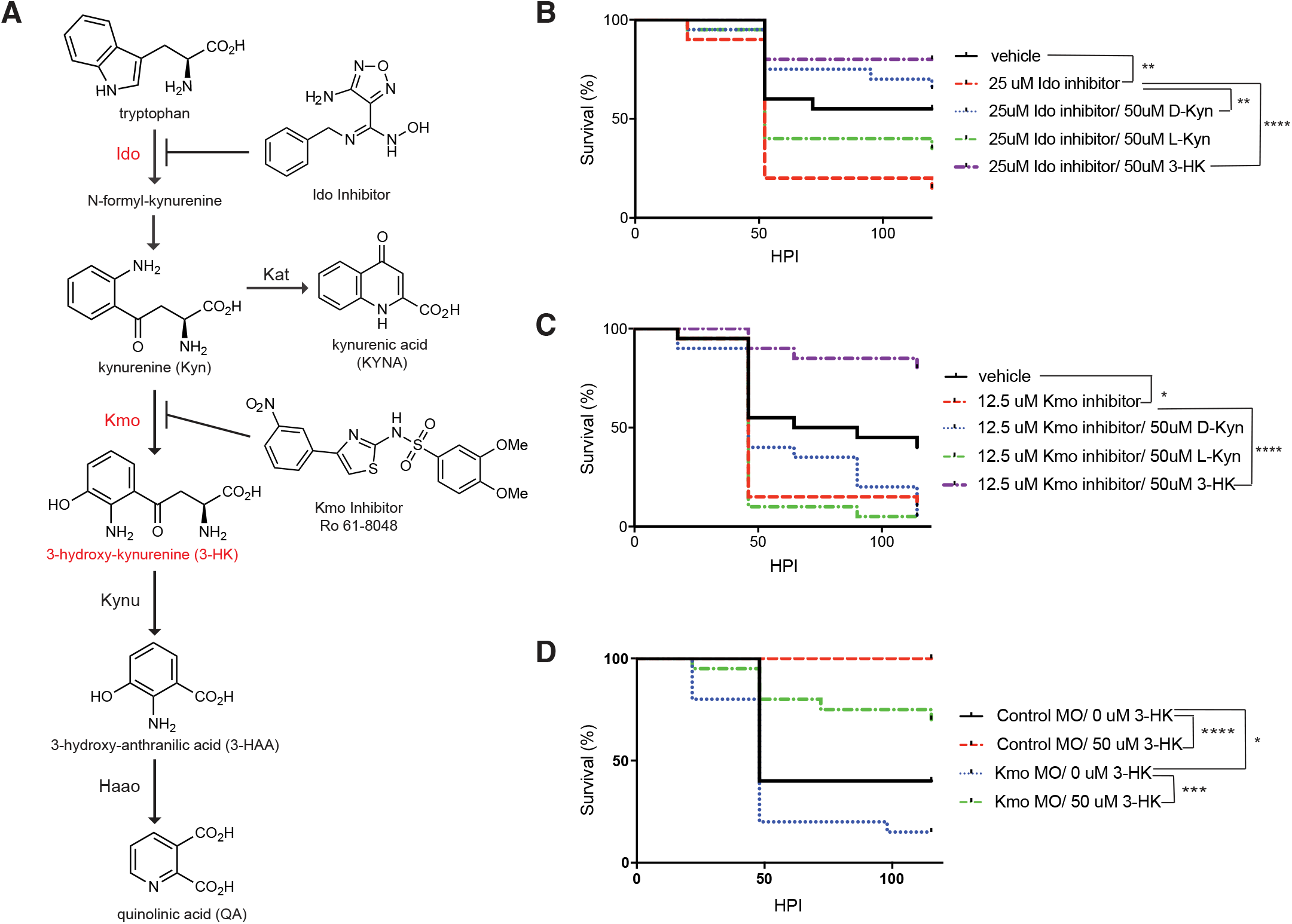
Endogenous production of 3-HK through the kynurenine pathway is important for defense against lethal *S.* Typhimurium infection. (**A**) Schematic representation of the kynurenine pathway. (**B**) 4-amino-N’-benzyl-N-hydroxy-1,2,5-oxadiazole-3-carboximidamide, an Ido inhibitor, sensitizes larvae to sublethal challenge with *S.* Typhimurium (275+/-71 CFU) and the sensitivity is reversed with D-Kynurenine or 3-HK (**, p=0.0049, Ido inhibitor; **, p=0.0013, D-Kynurenine; ****, p<0.0001). (**C**) Ro 61-8048, a Kmo inhibitor, sensitizes larvae to sublethal challenge with *S.* Typhimurium (329+/-62 CFU) and the sensitivity is significantly reversed with 3-HK (*, p=0.0289; ****, p<0.0001). (**D**) Kmo morphants are sensitized to sublethal challenge with *S.* Typhimurium (385+/-98 CFU) and the sensitivity is significantly reversed with 3-HK (*, p=0.0248; ***, p=0.0003). (**B**,**C**,**D**) For each graph, N=20 larvae/ condition; statistical significance was determined with the log rank test; data are representative of at least 3 independent experiments.

Given the benefit of exogenous 3-HK in promoting larval survival from lethal *S.* Typhimurium challenge, we sought to test whether the endogenous production of 3-HK by the host kynurenine pathway plays a role in defense against infection. We treated infected larvae with small molecule inhibitors of Ido (*19*) and Kmo (*20*) to prevent the catabolism of tryptophan to kynurenine and then to 3-HK (Fig. 2A), thus blocking endogenous 3-HK production.

Inhibition of both Ido and Kmo sensitized larvae to sublethal challenge with *S.* Typhimurium; importantly, this sensitivity was reversed with the addition of exogenous 3-HK (Fig. 2B and C). Further, Ido inhibitor induced sensitivity to *S.* Typhimurium infection could be reversed with exogenous addition of kynurenine, a product produced downstream of Ido, with D-kynurenine providing more significant rescue than the L-enantiomer (Fig. 2B). In contrast, Kmo inhibitor-induced sensitivity to *S.* Typhimurium infection could not be significantly reversed with either L-or D-kynurenine (Fig. 2C). Finally, genetic depletion of *kmo* using a morpholino resulted in sensitization to infection which could be reversed with exogenous 3-HK, thus phenocopying the effect of the Kmo inhibitor (Fig. 2D). Taken together, these results reveal that the host kynurenine pathway, and in particular, Ido and Kmo activity, are important for control of systemic infection in zebrafish and specifically, that endogenous production of 3-HK is part of the innate immune response that mediates defense against infection.

### 3-HK treatment does not enhance survival via known immune mechanisms involving the kynurenine pathway

The kynurenine pathway has been described to play a role in vertebrate immunity by a variety of different mechanisms involving different metabolites in this pathway (*8, 21*). We thus started by examining the ability of exogenous addition of other known kynurenine pathway metabolites to promote survival following infection and found that 3-HK was unique in its potent activity in promoting survival; tryptophan, L-and D-kynurenine, 3-HAA, QA, kynurenic acid (KYNA) and xanthurenic acid were all inactive (fig. S7B-E), despite confirmation of their ability to penetrate fish tissue based on their ability to affect other phenotypes (Fig. 2B and fig. S7F and G).

3-HK and the downstream metabolite 3-HAA have been reported to act as pro-oxidants (*22–24*) or anti-oxidants (*25, 26*), depending on the environment. We thus measured ROS levels in both infected and uninfected fish with and without 3-HK. We found that ROS levels were in fact lower in 3-HK treated fish (fig. S8A), consistent with 3-HK’s ability to act as an anti-oxidant (*25, 26*). Of note, both 3-HK and 3-HAA had anti-oxidant activity in this setting (fig. S8A and B). However, since only 3-HK but not 3-HAA improved survival, 3-HK’s anti-oxidant activity in this setting alone cannot explain the survival benefit (Fig. 1B; fig. S7D).

Other mechanisms by which the kynurenine pathway is known to impact the host (*8, 21*) also do not explain the observed 3-HK survival effect in a straightforward way. First, Idomediated depletion of intracellular tryptophan has been described to control replication of tryptophan auxotrophic pathogens (*27–30*); one could postulate that 3-HK might induce a feedback mechanism resulting in tryptophan depletion, similar to what has been described for kynurenine (*31*). However, neither the *Pseudomonas* nor the *Salmonella* strain used is a tryptophan auxotroph, exogenous tryptophan supplementation has no effect on survival (fig. S7B), and Ido-inhibitor induced sensitivity is reversed with the addition of both kynurenine and 3-HK (Fig. 2B). Alternatively, several metabolites such as kynurenine, kynurenic acid (KYNA; Fig. 2A) and xanthurenic acid (XA) (fig. S7A) are known to agonize the aryl hydrocarbon receptor (AHR) to impact the function of immune cells, particularly T cells (*e.g*., Treg differentiation)(*8, 21, 32*). However, T cells are absent in zebrafish larvae at the developmental stage utilized (*3*) and treatment with the AHR direct agonists kynurenine, KYNA or XA (fig. S7C and E) or the AHR antagonist CH-223191 (fig. S8C)(*33*) had no significant effect on survival. Finally, while the kynurenine pathway ultimately culminates in NAD+ synthesis, we found that 3-HK immersion did not increase NAD+ levels as a means to promote host survival after infection (fig. S8D). Thus, none of these mechanisms by which Ido and kynurenine metabolites are known to impact the host satisfactorily explain the observed 3-HK survival benefit.

### Expression analysis of the *S.* Typhimurium-macrophage interaction in the presence and absence of 3-HK

Given that 3-HK’s survival benefit unveiled a novel mechanism for the host kynurenine pathway in immunity with 3-HK being the specific metabolite promoting macrophage control of intracellular bacteria, we looked further into 3-HK’s effect on macrophages during infection. We performed bulk expression analysis of macrophages in infected fish in the presence and absence of 3-HK at 4 hpi to characterize the *S.* Typhimurium-macrophage interaction. We isolated mCherry positive macrophages from *mpeg1:mCherry* larvae infected with *S.* Typhimurium expressing GFP. Sorting based on their GFP fluorescence allowed us to distinguish subpopulations of invaded (GFP^+^) and exposed (GFP^-^; not invaded) macrophages from infected fish as well as naïve, resting macrophages from uninfected fish that had been treated with 3-HK or with vehicle control (Fig. 3A).

**Fig. 3.**
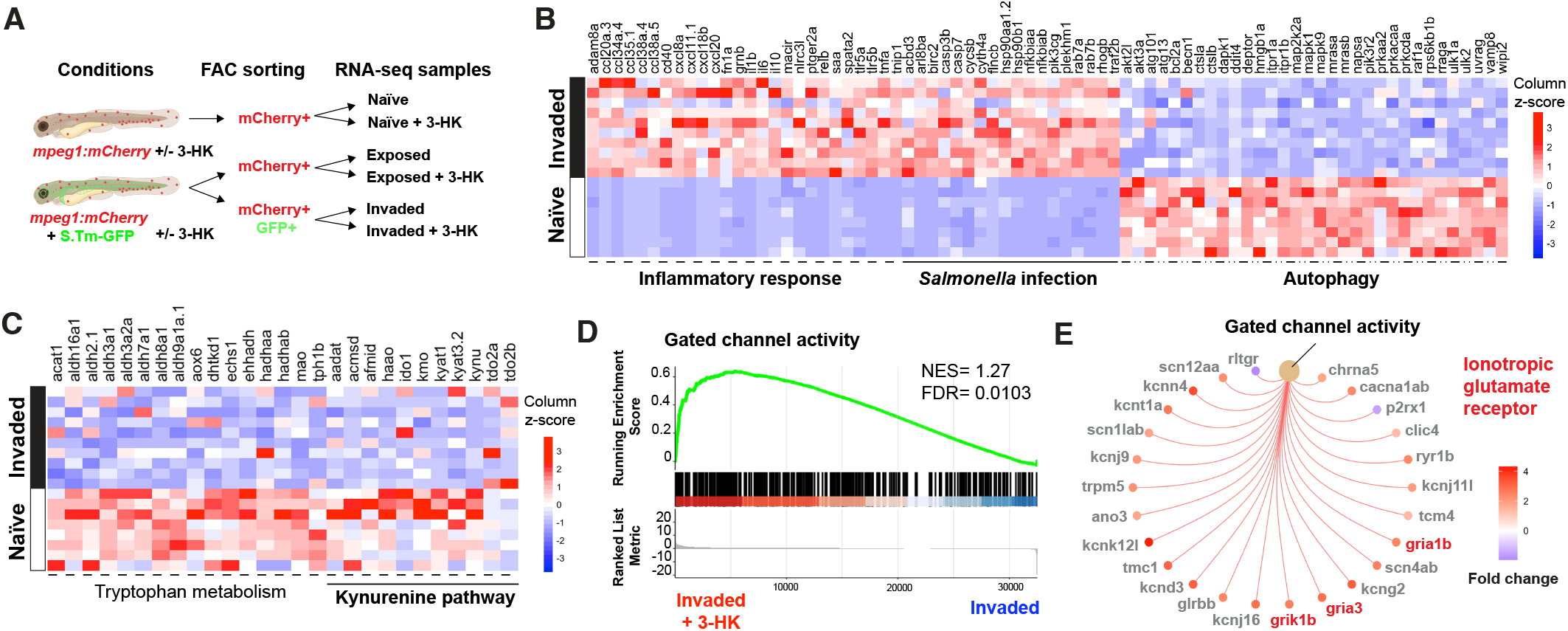
*S.* Typhimurium invasion of macrophages suppresses tryptophan kynurenine pathway metabolism and 3-HK supplementation upregulates the expression of genes associated to gated ion channels activity including that of ionotropic glutamate receptors. (**A**) Ilustration of bulk RNA-seq analysis from sorted populations of macrophages isolated from uninfected and *S.* Typhimurium infected animals with and without 3-HK treatment at 4 hpi. Briefly, 72 hpf embryos were infected or uninfected, 3-HK treated or vehicle treated, and at 4 hpi a single cell suspension of groups of 5 embryos per condition was prepared for fluorescence activated cell sorting according to fluorescence emission (*i.e.,* mCherry+ or mCherry+/GFP+). A range of 8-13 biological replicates/condition were collected from 2 independent infection experiments (837.5 +/-117.5 CFU and 770 +/-45 CFU). (**B**-**C**) Heatmaps showing selected genes differential expression from significant biological processes identified by Gene Set Enrichment Analysis in *S.* Typhimurium invaded versus naïve macrophages. (**B**) Gene sets belonging to pro-inflammatory responses, *Salmonella* infection, autophagy and (**C**) tryptophan metabolism genes are shown and clustered according to category (FDR < 0.05 for all 4 pathways). *S.* Typhimurium invasion of macrophages upregulates genes related to pro-inflammatory responses and *Salmonella* infection, and supresses the antimicrobial autophagy response. (**D**) Gene Set Enrichment Analysis plot of gene sets related to gated channel activity in *S.* Typhimurium invaded macrophages from 3-HK treated animals versus untreated animals. 3-HK treatment upregulates the expression of gated ion channels in *S.* Typhimurium invaded macrophages. (**E**) Cnet plot of leading edge genes associated with gated channel activity from the Gene Set Enrichment Analysis in (**D**) highlighing in red genes encoding ionotropic glutamate receptors of the AMPA (*gria1b*, *gria3*) and Kainate (*grik1b*) receptors subclasses (padj < 0.05 for all genes displayed). NES: normalized enrichment score; FDR: false discovery rate.

We first conducted differential expression analysis between *S.* Typhimurium invaded macrophages from infected fish and naïve macrophages isolated from control, uninfected fish. As expected, functional analysis by gene set enrichment analysis (GSEA) of differentially expressed genes revealed upregulation of canonical pathways associated with inflammation including the prototypical pro-inflammatory response genes *tnf-a, il-1b, il-6.* We also found upregulation of gene sets specifically known to be related to *Salmonella* infection including genes associated with intracellular vacuolar replication *plekhm1*, *rab7a, rab7b* and cell death *casp7a, casp7b* and *traf2b*(*34–37*) (Fig. 3B and fig. S9A and B). In addition, we found that *S.* Typhimurium invasion of macrophages suppressed expression of the antimicrobial autophagy response, including genes encoding for the evolutionarily conserved autophagy-related proteins (ATGs) that control dynamic membrane events during autophagosome biogenesis, *becn1(ATG6)*, *atg13* and *atg101* (Fig. 3B and fig. S9C)(*38*). Autophagy has been shown to play a role in *S.* Typhimurium infection in mammals and zebrafish larvae (*15, 39, 40*).

We then compared the expression programs between *S.* Typhimurium invaded and exposed macrophages from infected fish, and then between exposed macrophages from infected fish and naïve macrophages from uninfected fish. Compared to exposed macrophages from infected fish, invaded macrophages upregulated genes involved in vacuole, endosome and lysosomal functions and regulation of intracellular pH (fig. S9D and E). When compared to naïve macrophages from uninfected fish, similar to invaded macrophages, exposed macrophages also upregulated pro-inflammatory response genes (fig. S9F). The fact that these results are largely as expected and consistent with what is known about *S*. Typhimurium infection confirmed the reliability and robustness of the transcriptional data.

Notably, GSEA comparison of *S.* Typhimurium invaded macrophages with exposed or naïve macrophages revealed that *S.* Typhimurium intracellular invasion suppresses tryptophan metabolism with significant downregulation of genes encoding enzymes of the kynurenine pathway including *kmo*, *kynu* and *haao* (Fig. 3C and fig. S9G). As previous reports have shown that IFN-γ production and upregulation in the kynurenine pathway are associated with protection from infection (*27, 28, 41–43*), these results are consistent with the idea that *S.* Typhimurium invasion downregulates the kynurenine pathway in macrophages to promote its own survival in some settings (*44*).

Finally, we compared the expression programs of 3-HK treatment versus vehicle control in *S.* Typhimurium invaded and exposed macrophages isolated from infected fish, and naïve macrophages isolated from uninfected fish. The majority of transcriptional changes induced by 3-HK treatment in each macrophage subpopulation were in several classes of ion channels including both voltage- and ligand-gated ion channels (Fig. 3D and E and fig. S9H and I). In particular, 3-HK treatment upregulated the expression of iGluR subunits of the KAR (*grik1b*) and α-amino-3-hydroxy-5-methyl-4-isoxazolepropionic acid (AMPAR; *gria1b* and *gria3*) receptor subclasses in *S.* Typhimurium invaded macrophages (Fig. 3E). Interestingly, the kynurenine metabolites QA and KYNA are known agonists and antagonists, respectively, of iGluRs of the NMDAR subclass (*4, 5, 7*) (Fig. 2A), with the endogenous balance of these metabolites associated with the progression of some neurodegenerative diseases and mood disorders (*5, 7, 8*). However, we did not observe that 3-HK induced differential regulation of NMDAR subunits (fig. S9J). Instead, 3-HK treatment induces transcriptional changes in iGluRs of the AMPA and KAR subclasses.

Collectively, these results reveal that *S.* Typhimurium infection in zebrafish larvae can lead to a suppression of the kynurenine pathway in *S.* Typhimurium invaded macrophages. Meanwhile, 3-HK supplementation overall induces changes in the expression of ion channels in macrophages independent of infection state, suggesting that 3-HK could modulate the activity of iGluR receptors of the AMPAR and KAR subclasses, as activity dependent differential transcriptional regulation has been noted for iGluRs (*45, 46*).

### 3-HK protects from infection through inhibition of iGluRs of the kainate class (KARs)

Taking a complementary method to gain insight into 3-HK’s mechanism of action, we utilized a similarity ensemble approach (SEA)(*47*) to predict possible receptors for 3-HK. SEA is a cheminformatic analysis that identifies protein targets based on the chemical structural similarity of a ligand to other known small molecule ligands with known, cognate receptors. In addition to kynureninase (KYNU), the enzyme that converts 3-HK to 3-HAA, and amino acid transporters, SEA predicted that 3-HK is a potential ligand of iGluRs of the KAR and AMPAR, but not NMDAR, subclasses. This prediction strikingly mirrored the observed transcriptional changes in *S.* Typhimurium invaded macrophages following 3-HK treatment and pointed to a potential role for KARs or AMPARs in modulating host response to infection.

KARs and AMPARs form two subclasses of iGluRs that are most highly expressed in neurons of both the central and peripheral nervous systems, where they are involved in synaptic transmission and plasticity, though they have been identified in other cell types including leukocytes, platelets and pancreatic islet cells (*48–51*). In mammals, these proteins are glutamate-gated ion channels that conduct cations upon ligand binding; they have also been reported to have metabotropic activity as well (*52, 53*). Both KARs and AMPARs are formed by tetramerization of several possible pore-forming subunits. KAR subunits are encoded by five different genes, *Grik1-Grik5*, and Neto auxiliary subunits (*52, 54*). *Grik1-Grik3* encode subunits that may either homo- or hetero-tetramerize, while *Grik4* and *Grik5* encode high-affinity subunits that may only hetero-tetramerize with subunits encoded by *Grik1*-*Grik3* (*52*). The genetic architecture of AMPARs is similarly complex as this sub-class of iGluRs are formed by homo- or hetero-tetramerization of four subunits encoded by 4 different genes, *Gria1-Gria4,* also associated with a variety of auxiliary subunits (*55, 56*).

Given the genetic complexity of KARs and AMPARs and their numerous combinatorial subunits, we took a chemical genetic approach, testing whether known ligands (agonists or antagonists) of KARs/AMPARs could recapitulate 3-HK’s survival benefit in infection. Indeed, two structurally distinct antagonists, the competitive KAR/AMPAR antagonist CNQX and the more specific non-competitive KAR antagonist NS 3763 (*57–59*) phenocopied 3-HK, providing significant protection from lethal infection (Fig. 4A and B). Of note, like 3-HK, neither molecule had any effect on bacterial growth in axenic culture (fig. S2D-G). In contrast, the KAR agonist, kainate, significantly sensitized larvae to sublethal challenge, while NS 3763 was able to compete out kainate-induced sensitivity (Fig. 4C). These results independently confirm the role of KARs in modulating survival.

**Fig. 4.**
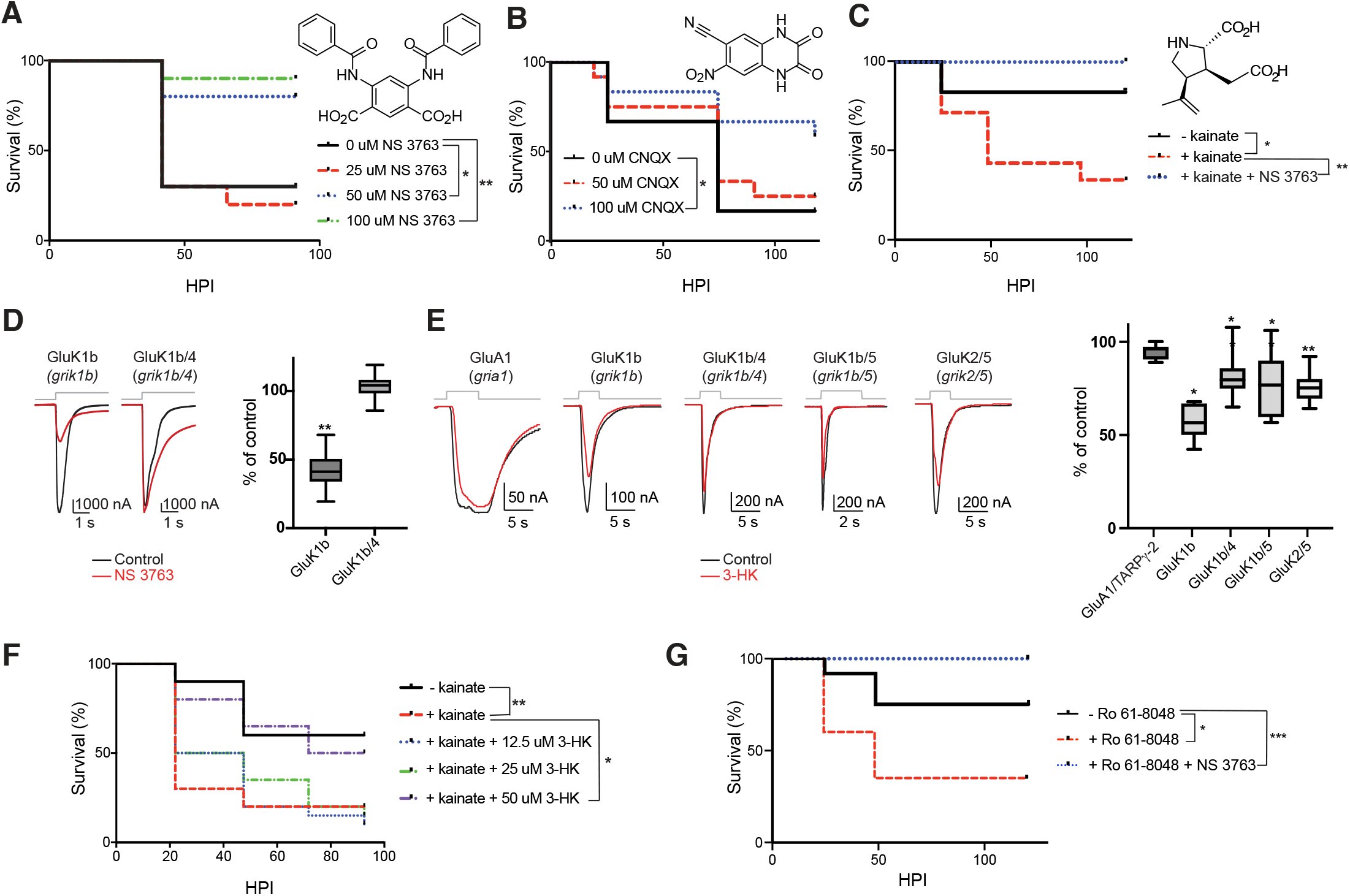
The KAR subunit encoded by *grik1b* mediates 3-HK’s survival effect during infection. Larvae (72 hpf; N=10-12/condition) were infected with *S.* Typhimurium ((**A**) 747+/-235 CFU; (**B**) 276+/-51 CFU) and treated with vehicle control (DMSO) or indicated concentrations of the non-competitive KAR antagonist, NS 3763 (**A**), or the competitive KAR antagonist, CNQX (**B**), both of which significantly promoted survival of infected larvae ((**A**) *, P=0.0285; **, P=0.0076; (**B**) *, P=0.05; log-rank test). (**C**) The KAR agonist, kainate (350uM), sensitized larvae to sublethal challenge with *S.* Typhimurium (250 +/-10 CFU) and the sensitivity was reversed with NS 3763 (50uM) (N=12 embryos/ condition; *, P=0.0517; **, P=0.0019; log rank test). (**D**) NS 3763 significantly inhibited glutamate-evoked currents through GluK1b (*grik1b*) but not GluK1b/4 (*grik1b/4*) coexpressed with Neto2 auxiliary subunit. Representative traces (left) and quantification of relative ratios of 100 uM glutamate-evoked currents with 50-75 uM NS 3763 for replicate measurements (right) are shown (N=8 oocytes/group; whiskers: min. to max; Wilcoxian signed rank test; **, P=0.0078 (GluK1b). (**E**) 3-HK significantly inhibited glutamate-evoked currents through GluK1b (*grik1b*), GluK1b/4 (*grik1b/4*), GluK1b/5 (*grik1b/5*) and GluK2/5 (*grik2/5*) coexpressed with Neto2 auxiliary subunit in comparison to a structurally similar AMPAR receptor (GluA1 (*gria1*) coexpressed with TARPγ-2 auxiliary subunit. Representative traces (left) and quantification of relative ratios of 100 uM glutamate-evoked currents with 50 uM 3-HK for replicate measurements (right) are shown (N=7-9 oocytes/group; whiskers: min. to max; Wilcoxian signed rank test; *, P=0.0156 (GluK1b), P=0.0234 (GluK1b/4), P=0.0195 (GluK1b/5); **, P=0.0078 (GluK2/5). (**F**) The KAR agonist, kainate (175uM), sensitized larvae to sublethal challenge with *S.* Typhimurium (361+/-34 CFU) and the sensitivity was reversed with 3-HK (N=20 embryos/ condition; *, P=0.0116; **, P=0.0014; log rank test). (**G**) The Kmo inhibitor Ro 61-8048 (25uM) sensitized larvae to sublethal challenge with *S.* Typhimurium (175 +/-5 CFU) and the sensitivity was reversed with NS 3763 (50uM) (N=12 embryos/ condition; *, P=0.0342; ***, P=0.0008; log rank test).

To explore the possibility that 3-HK, like NS 3763, is an antagonist of KARs, we measured its ability to inhibit glutamate-evoked currents in *Xenopus laevis* oocytes with heterologously expressed KAR subunits. Since NS 3763 is selective for mammalian *Grik1-* encoded subunits (*57, 59, 60*) and *grik1b* was the only KAR subunit upregulated in macrophages of 3-HK treated, infected zebrafish (Fig 3E), we focused on the two zebrafish paralogs of *Grik1*, *grik1a* and *grik1b*. We heterologously expressed full-length *grik1a* and *grik1b* as well as that of the high-affinity subunits encoded by *grik4* and *grik5* from zebrafish in *Xenopus laevis* oocytes by co-injecting cRNAs of corresponding KAR subunits and Neto2 auxiliary subunit. We were unable to evaluate channels encoded by *grik1a* because they did not show activity in cRNA-injected oocytes (fig. S10A). We confirmed that NS 3763 inhibited the peak amplitude of glutamate-evoked currents of *Danio rerio grik1b-*encoding GluK1b homo-tetramers, but not heteromers encoded by *grik1b and grik4* and seemed to slow the current decay of both channels (Fig. 4D). This subtype specificity resembles that in mammalian encoded KARs reported previously (*57, 59, 60*). 3-HK also significantly inhibited currents from oocytes expressing channels encoded by zebrafish *grik1b*, with the greatest inhibition of homo-tetramer channels encoded by *grik1b* (Fig. 4E), similar to NS 3763, and to a lesser extent hetero-tetramer channels encoded by zebrafish *grik1b/grik4* and zebrafish *grik1b/grik5* (Fig. 4E; 55-80% inhibition). 3-HK did not inhibit the structurally similar AMPAR encoded by *Gria1* and *TARP γ-2* auxiliary subunit (Fig. 4E and fig. S10B and C). Importantly, when we tested the ability of 3-HK to complete away kainate-induced sensitivity to infection in zebrafish, indeed, 3-HK, like NS 3763, restored survival, consistent with its activity targeting KARs (Fig. 4F). 3-HK’s inhibition of glutamate-evoked currents, taken together with its ability to reverse kainate-induced sensitization to infection, demonstrate that exogenous 3-HK activity at KARs can account for its survival benefit.

Finally, we sought to determine if endogenous production of 3-HK via the kynurenine pathway also promotes zebrafish survival to infection through KARs. We tested whether the KAR antagonist NS 3763 could rescue the sensitivity to infection induced by inhibition of endogenous 3-HK production via Kmo chemical inhibition by Ro 61-8048. Indeed, NS 3763 treatment reversed the increased sensitivity to infection due to Kmo inhibition, protecting fish from lethal infection (Fig. 4G). Thus, KAR antagonism reversed the loss of endogenous 3-HK during infection, demonstrating an interaction between the endogenous production of 3-HK by the kynurenine pathway and KAR channels.

## DISCUSSION

In a whole organism screen for small molecules that promote zebrafish embryo survival following lethal infection, we found that the tryptophan catabolite 3-HK confers a profound survival advantage to zebrafish larvae infected with *P. aeruginosa* and *S.* Typhimurium. It does not act like a typical antibiotic as it does not have any direct antibacterial activity *in vitro* but does promote macrophage-dependent control of bacterial expansion *in vivo*. Moreover, endogenous host production of 3-HK via the kynurenine pathway is required for defense against *S.* Typhimurium infection as chemical inhibition or genetic knock-down of Kmo sensitizes animals to infection. Two complementary approaches, transcriptional profiling of infected macrophages in response to 3-HK and SEA (*47*), a computational analysis that identifies putative protein targets based on ligand structural similarities, both converged on a role for KARs or AMPARs as possible receptors for 3-HK.

We independently implicated KARs in infection through a series of chemical genetic experiments in which two structurally distinct antagonists, one of which is highly specific for *Grik1*-encoded channels (NS 3763) (*57, 59, 60*), provided significant protection from infection. Conversely and importantly, treatment with the KAR agonist kainate, sensitized fish to infection which could be competed away with NS 3763. Thus, KARs in and of themselves play a role in modulating host responses to S. Typhimurium infection.

3-HK’s modulation of KAR activity was demonstrated electrophysiologically in *Xenopus* oocytes. Importantly, this activity at KARs accounts for 3-HK’s survival benefit during infection as 3-HK phenocopies the known KAR antagonist NS 3763 in its rescue of infected zebrafish from sensitization induced by the KAR agonist kainate. Finally, we demonstrated an intersection between the endogenous production of 3-HK by the host kynurenine pathway and KAR antagonism in immunity, as the loss of endogenous 3-HK is complemented by antagonizing KARs with NS 3763. The implication is that *grik1*-encoded KAR channels are a target of endogenously produced 3-HK and that this interaction is important in determining the outcome of infection. It also raises the possibility that this interaction may play a role in other settings as well (*e.g.*, neurodegeneration).

This infection model and the identification of 3-HK in a whole organism chemical screen for modulators of survival enabled the discovery that the kynurenine pathway regulates innate immunity through a novel mechanism involving 3-HK activity at KARs. In mammals, *Ido1* and other kynurenine pathway enzymes, including *Kmo* (*8, 27, 28, 41, 42*) are induced by IFN-γ and other inflammatory stimuli. However, *Ido* and *Kmo* have been noted to be downregulated following infection (*44*) when IFN-γ signaling is blocked (*e.g.*, in IFN-γR knock-out mice). In zebrafish larvae, the downregulation of *ido* and *kmo* in invaded macrophages is similar to this latter finding (*44*), consistent with previous observations that zebrafish IFN-γ orthologues are not upregulated in response to infection at early developmental stages (*61*). This downregulation is also consistent with the emerging concept that *S.* Typhimurium can reprogram and co-opt macrophage metabolism to promote its own intracellular survival and replication (*62, 63*). Thus, the zebrafish larval host model, where kynurenine pathway enzymes are downregulated following infection, unmasked a novel mechanism by which the specific metabolite 3-HK contributes to host defense. The finding that 3-HK is specific to gram-negative organisms like *S.* Typhimurium and *P. aeruginosa* as opposed to organisms like *S. aureus* or *M. marinum*, might suggest this mechanism may be important when the host is sensing and responding to pathogen associated molecular patterns common to gram-negative organisms, such as LPS.

The kynurenine pathway has been implicated in the control of a variety of pathogens including bacteria, viruses, and parasites. For some obligate intracellular tryptophan auxotrophs including *T. gondii* and some Chlamydial species, IDO-induced tryptophan deprivation is the primary mechanism by which the kynurenine pathway exerts control over pathogen burden as restoration of tryptophan prototrophy in these organisms relieves this control (*27–29, 64*). For other pathogens that are not obvious tryptophan auxotrophs, however, the mechanism of control remains poorly understood, though it has been attributed to toxic effects of metabolites downstream of IDO1 (*42–44, 65*). Here we find that 3-HK can functionally antagonize KARs to uniquely promote host survival in the setting of *S.* Typhimurium infection in zebrafish, thereby elucidating a novel mechanism by which this downstream metabolite can promote defense against infection. This mechanism may be relevant to some of the other infections where the mechanism has to date remained elusive.

3-HK antagonism of KARs could promote control of bacterial burden and host survival directly or indirectly in macrophages. While iGluRs are known to be expressed on immune cells (*48, 50*), including in fish macrophages (Fig. 3E), if or how they act to support the function of these cell types *in vivo* remains relatively poorly understood (*48, 50*). Thus, whether the clear induction of neurotransmitter receptors, including KARs, in macrophages of 3-HK treated fish is indicative of 3-HK’s direct activity on macrophages to control bacterial burden or is irrelevant, merely echoing transcriptional changes which are actually occurring and more relevant in an alternative cell type, including possibly neurons, remains to be determined.

While this work newly links the kynurenine pathway and iGluRs in host defense to bacterial infection, the relationship between this pathway and receptor family has been well established in the context of neurotransmission and neurodegeneration. Inflammatory stimuli can activate the kynurenine pathway in the CNS, including in microglia, as well as in the periphery (*5, 8*). Within the CNS, this activation can result in synthesis of metabolites including QA (*5, 8*), which can in turn stimulate NMDARs expressed in neighboring neurons and contribute to neuronal death in some neurodegenerative disorders, an effect that can be potentiated by 3-HK (*66*). 3-HK itself is toxic to particular subsets of neurons, an activity attributed to its promotion of ROS (*22–24*). For this reason, therapeutic efforts have focused on inhibiting KMO, a critical branchpoint in the pathway, to decrease the availability of the NMDAR agonist QA and neurotoxic metabolite 3-HK, while driving kynurenine metabolism toward the neuroprotective metabolite KYNA (*5, 7, 8*), a potent antagonist of NMDARs with much weaker antagonist activity at other iGluRs (*67–70*). This work now expands the relationship between the kynurenine pathway and iGluRs, linking an additional metabolite (3-HK) and additional iGluR subtypes (KARs) in a novel way in support of a new function: immune defense to bacterial infection.

Beyond its impact on host defense, the discovery of 3-HK’s antagonism of KARs also has potentially significant implications for neurotransmission and neuronal cell death. It adds a layer of complexity in interpreting how these metabolites may be affecting synaptic transmission in the CNS or the periphery where KARs are expressed. While it is expected that QA will agonize NMDARs in neighboring neurons to increase neuronal excitability, the effect of KAR antagonism is more complicated. KARs are the least well understood among iGluRs but are believed to play more of a modulatory role in regulating synaptic transmission, functioning post-synaptically in regulating neuronal excitability and pre-synaptically by modulating the release of neurotransmitters such as GABA and glutamate (*52, 71*). Thus, they function to tune the threshold of neurotransmitter release in neurons where they are expressed, resulting in either activation or inhibition (*52, 71*). Their role *in vivo* can be further complicated by the fact that in addition to functioning as conventional ligand-gated ion channels, KARs have also been shown to signal metabotropically (*52, 53, 71*). Finally, predicting how 3-HK activity at KARs might contribute to neuronal activation or inhibition is also likely to be dependent on the neuronal cell type in which KARs are expressed as well as the local environment of other kynurenine metabolites, neurotransmitters, and immune cell-types present. Despite the fact that there is still much to be understood, this work demonstrates that 3-HK activity at KARs can have a profound effect on organism physiology in settings where the kynurenine pathway is active, such as during infection or where its production under conditions of ‘sterile’ inflammation is detrimental to host health (*72*) including acute pancreatitis (*73*) or the neuroinflammation in Huntington’s and/ or Alzheimer’s disease (*5*).

Taken together, this work reveals a new mechanism by which tryptophan metabolism modulates KARs to regulate host defense to bacterial infection. It also broadens the significance of the relationship between the kynurenine pathway and iGluRs beyond neurotransmission to host defense and increases the complexity by which kynurenine metabolites interact with iGluRs to regulate inflammatory states. The potentially wide impact of 3-HK’s activity at KARs, coupled with recent interest in kynurenine pathway enzymes as therapeutic targets in conditions ranging from inflammation, neurodegeneration, and cancer (*5, 7, 8*), highlights the need for a greater understanding of the role of the interaction between the kynurenine pathway and KARs in immune responses, neurodegeneration, and other inflammatory disorders more generally.

## Acknowledgments

We thank S. Shaw for contributing the Prestwick collection bioactive library for the chemical screen and F. Ausubel, R. Xavier, and S. Walker for contributing bacterial strains. We thank L. Dong and the Harvard Chan Bioinformatics Core for their help with their freely available resources and tools for next generation sequencing data analysis utilized in this study. We also thank N. Hacohen and T. Eisenhaure for providing the in-house adaptation of the Smart-seq2 protocol for bulk immune cell RNA-seq used in this study. We also thank Maggie Ma and Erica Ueda for their assistance pinning compounds. Schematic image in Figure 3 was created with BioRender.com.

## Author contributions

Conceptualization: MPK, AEC, DTH

Methodology: MPK, AEC, DTH, ST

Investigation: MPK, AEC, EJS, CC, ERG, SC, JSWL, SMB, CRO

Visualization: MPK, AEC, EJS, CC, ERG, SMB

Funding acquisition: MPK, DTH, ST, EJS

Project administration: AEC, DTH, ST Supervision: DTH, ST, AEC

Writing – original draft: MPK, AEC, DTH

Writing – review & editing: MPK, AEC, DTH, ST

## Data and materials availability

All other data are available in the manuscript or the supplemental information.

## Supplemental Information

### Materials and Methods

#### Bacterial Strains and growth conditions

The following strains were used to infect zebrafish embryos and larvae: *Salmonella enterica* serovar Typhimurium SL1344 (gift from Ramnik Xavier, MGH), *S.* Typhimurium-GFP (*74*)*, Pseudomonas aeruginosa* PA14*1′oprM*, *Staphylococcus aureus* Newman (gift from Suzanne Walker, HMS), and *Mycobacterium marinum/GFP*(*75*). The *S.* Typhimurium inoculum was prepared by inoculating an overnight culture grown in LB at 37°C with shaking. The overnight culture was sub-cultured (1:500) and grown for ∼3 hours at 37°C with shaking until mid-log phase was reached. Bacteria were then pelleted by centrifugation, washed once in 1X PBS and resuspended at the desired inoculum size in 1X PBS and 0.05% phenol red, to facilitate visualization of injecting the inoculum. For some experiments, following washing in 1X PBS, bacteria were resuspended at 4X the desired inoculum size in 1X PBS with 20% glycerol and frozen back in single use aliquots at −80C. Individual aliquots were then thawed, diluted in 1X PBS and 0.05% phenol red for direct inoculation. No differences in infection were noted between bacterial inoculates prepared by either method. The bacterial inoculum for infections with *P. aeruginosa* PA14*1′oprM* and *M. marinum*/GFP were prepared as previously described(*9, 75, 76*). The *S. aureus* inoculum was prepared by sub-culturing an overnight culture in tryptic soy broth (TSB) at 37°C with shaking for 3 hours. Bacteria were then pelleted by centrifugation and resuspended in PBS with 0.05% phenol red for injection into zebrafish embryos.

#### In vitro bacterial growth inhibition assays

*S.* Typhimurium was grown overnight at 37°C with shaking and then sub-cultured (1:1000) in 50 ml LB +/-indicated concentrations of 3-HK and incubated at 37°C with shaking over 24 hours. The OD600 of each culture was measured over 24 hours by cuvette on a Spectromax M5 spectrophotometer (Molecular Devices). For *in vitro* experiments with *S.* Typhimurium and *P. aeruginosa* growth in LB and M9, bacteria were grown over night in LB at 37°C with shaking. Saturated overnight cultures were sub-cultured and grown at 37°C with shaking until log phase, diluted to an OD600 of 0.05 in either LB or M9 and arrayed in 96-well format with two-fold serial dilutions of 3-HK, NS 3763 or CNQX ranging from 0-250 uM. The plate was incubated in a humidity chamber at 37°C and the OD600 measured every 15 minutes with 60 seconds shaking before each reading over a ∼24 hours using a Tecan Spark 10M plate reader (Tecan).

#### Zebrafish Lines

Wildtype zebrafish from the AB line and 2 transgenic lines, Tg(*mpeg1:mCherry*) (*gl22Tg*) with mCherry-labeled macrophages under the *mpeg1* (*macrophage expressed gene 1*) promoter(*77*) and Tg(*mpx:EGFP)^i114^*with EGFP-labeled neutrophils under the *mpx* (*myeloperoxidase*) promoter(*78*), were used in this study. Zebrafish were maintained in compliance with the guidelines and IACUC protocol approved by the MGH’s Institutional Animal Care and Use Committee.

#### Zebrafish infections and inoculum determinations

Zebrafish infections were carried out as previously described (*9, 75, 76*). Briefly, embryos (staged 50hpf) or larvae (72 hpf) were inoculated intravenously by microinjection into the duct of Cuvier/ common cardinal vein. Infected embryos/ larvae were then immediately immersed in embryo media (E3) containing vehicle control (DMSO or H2O) or small molecules of interest and incubated at 29°C. Thus, treatment of embryos and larvae with small molecules of interest began immediately following infection. Embryos/larvae were observed every day for survival with the scoring of living versus dead animals ascertained by the presence of a heartbeat and circulating blood under a stereomicroscope. The inoculum for each infection was retrospectively determined by inoculating tubes of PBS from the injection needle prior to and following a given set of injections and plating the inoculated tubes of PBS to LB agar (*P. aeruginosa* and *S.* Typhimurium), tryptic soy agar (*S. aureus*) or 7H10 agar (*M. marinum/*GFP) plates and counting the colonies that arose on each plate. Alternatively, embryos or larvae were homogenized immediately following inoculation and the homogenate plated to LB-streptomycin (25ug/ml; *S.* Typhimurium infections only) for CFU determinations as described below. Either method of inoculum determination provided comparable results.

#### Enumeration of infecting bacteria

Infected embryos/ larvae were euthanized with excess tricaine and incubated on ice for 5 min. Individual embryos/ larvae were then washed once in 1X PBS, resuspend in 0.1% Triton X-100 in 1X PBS and homogenized with sterile 5-mm stainless steel beads in three 2-min cycles of 40 Hz using a TissueLyser bead mill (Qiagen). Homogenates were vortexed well between bead mill cycles and just prior to plating the homogenate to LB-streptomycin (25ug/ml) agar plates for CFU determinations.

#### Chemical screen

The Prestwick Chemical library (Prestwick Chemical, 2 mg/mL stock concentrations in DMSO) of known bioactives and off-patent drugs was screened for small molecules that rescued zebrafish embryos from lethal challenge with *P. aeruginosa* PA14*1′oprM.* Since *P. aeruginosa* has a higher intrinsic resistance to multiple antibiotics due to its expression of multiple efflux pump systems, we sought to reduce its ability to efflux small molecules that might be hits in our screen by deleting the *oprM* gene, which encodes an outer membrane protein utilized by multiple efflux systems in *P. aeruginosa*. The resulting strain was more sensitive to antibiotics than the parent strain, PA14, and thus allowed us to screen the compound collection at a lower concentration than would have been possible with the parent strain (PA14) and still hit known antibiotics like ciprofloxacin within the collection. We first pre-screened 1056 compounds from the collection for toxicity by pinning 1 ul of compound/ well in a final volume of 200ul E3 media and 1% DMSO in 96-well format; wells containing E3 with 1% DMSO were included as vehicle controls for the toxicity screen. Four embryos (50 hpf) were arrayed per well and scored for viability, as determined by the presence of a heartbeat and circulating blood, over a period of 4 days. We found that 10.8% (114/1056) of the compounds tested were toxic to embryos over this length of time at this screening concentration and were subsequently omitted from the survival screen. To screen for compounds that promoted survival of infected fish, 1ul of each compound was pinned into 110ul E3 media in 96-well plate format and then 100ul from each of two compound wells were multiplexed manually, omitting compounds found to be toxic in the toxicity screen. Thus, each well for the screen contained 2 compounds in a final volume of 200ul E3 with ∼1% DMSO. Embryos (50 hpf) were inoculated intravenously by microinjection with an average of 2600 CFU *P. aeruginosa* PA14*1′oprM* and infected embryos were arrayed at 4 embryos/well in each screening plate immediately following infection. In general, embryos derived from a single clutch were arrayed evenly among all wells of a plate so that any clutch to clutch genetic diversity was evenly distributed among all wells for the screen. Each plate included 2 negative control wells (E3 with 1% DMSO) and 2 positive control wells (E3 with 12.5ug/ml ciprofloxacin in 1% DMSO); the Z’ factor for the assay was 0.56. Infected embryos were monitored over 4 days for the presence of a heartbeat and circulating blood under a stereo microscope. Any well having ≧1 surviving embryo 3 days post infection was considered to contain a hit. A total of 932 compounds were screened for survival with 67 wells determined to contain a hit; 21 of these 67 hit wells contained a known antibiotic and were not examined further. Compounds from the remaining 46 hit wells were repurchased and tested singly in three point dose response to determine efficacy in promoting zebrafish embryo survival.

#### Small molecules

3-hydroxy-D,L-kynurenine (purified stereoisomers of 3-HK were not available commercially) was purchased from Millipore Sigma, Biosynth Carbosynth or BOC Sciences. D-kynurenine, L-kynurenine, L-tryptophan, xanthurenic acid (XA), kynurenic acid (KYNA), 3-hydroxyanthranilic acid (3-HAA), quinolinic acid (QA), and β-nicotinamide mononucleotide (β-NMN) were purchased from Millipore Sigma. The Ido inhibitor, 4-amino-*N*’-benzyl-*N*-hydroxy-1,2,5-oxadiazole-3-carboximidamide, was purchased from Chembridge. The Kmo inhibitor,Ro 61-8048 and CNQX were purchased from Tocris. NS 3763 was purchased from Tocris, BioVision and Chembridge. Kainate was purchased from Tocris.

#### Zebrafish macrophage depletion experiments

Zebrafish embryos (2 dpf) were inoculated intravenously by microinjection into the duct of Cuvier/common cardinal vein with 1-3 nL of clodrosomes (2.5 mg/ml)(*79*). Selective elimination of macrophages and not neutrophils was confirmed by injection of liposomes and clodrosomes into double transgenic animals Tg(*mpeg1:mCherry/mpx:EGFP*) and immune cell quantification was conducted at 3 dpf by epifluorescence microscopy (Nikon 90i) imaging and manual quantification using Fiji(81).

#### Genetic knock-down experiments

Morpholinos (Gene Tools) were suspended in 1 mM sterile water, diluted to indicated concentrations and 1-2 nL was microinjected into 1-4 cell stage embryos. For knockdown of *pu.1,* a previously validated gene-specific morpholino was used; MO-pu.1 (1 nL, 0.5 mM; 5’-GATATACTGATACTCCATTGGTGGT-3’)(*13*). A custom morpholino was used to knockdown *kmo (kmo* MO: 2 nL, 125 uM morpholino; 5’-GATGTGAGAAAGCTGTCTCCATGTT-3’). A random control oligo 25-N, was used as a negative control (1-2 nL, 60-250 uM).

#### Ionotropic glutamate receptor activity

For expression in *Xenopus laevis* oocytes, full length *grik1b*, *grik1a*, *grik4* and *grik5* transcripts were amplified from zebrafish cDNA and cloned into 5’ entry vector pCR8/GW/TOPO (Invitrogen). Briefly, full length *grik1b*, *grik1a* and *grik5* were amplified from cDNA synthesized from RNA isolated from wildtype 3 dpf larvae with primers SC42 and AC936 (*grik1b*), SC8 and AC935 (*grik1a*), and CC189 and CC190 (*grik5*) using Q5 DNA polymerase (NEB). PCR products were 3’ A-tailed with Taq polymerase prior to Topo cloning into pCR8/GW/TOPO to create plasmids pCR8/gw/topo-*grik1b*, pCR8/gw/topo-*grik1a*X2, and pCR8/gw/topo-*grik5*. Full length *grik4* was amplified from cDNA as described above using primers SC179 and SC180 and assembled into the pCR8/GW/TOPO backbone using NEBuilder HiFi DNA Assembly Master Mix (NEB) and EcoRI digested and gel purified pCR8/gw/topo-*grik1b* (EcoRI digestion removes the *grik1b* insert leaving only vector backbone) to create pSC*grik4*. Plasmids pCR8/gw/topo-*grik1b*, pCR8/gw/topo-*grik1a*X2, pCR8/gw/topo-*grik5* and pSCgrik4 were used as middle entry vectors for Gateway (Invitrogen) cloning into pGEM-HE.

Two-electrode voltage clamp (TEVC) recording of *X. laevis* oocytes were performed as described(*80*). Briefly, cRNAs of KAR, AMPAR and auxiliary subunit constructs were subcloned into pGEM–HE vector and cRNAs were transcribed *in vitro* using T7 mMessage mMachine (Ambion) and corresponded cRNAs were co-injected into oocytes (0.1 ng KAR, 0.5 ng Neto2, 0.5 ng GluA1, 0.5 ng TARPγ-2). TEVC analysis was performed 3-5 days after injection at room temperature in recording solution containing (in mM): 100 NaCl, 1.0 KCl, 1.0 MgCl_2_, 0.5 BaCl_2_ and 5 HEPES (pH 7.4). The membrane potential was held at –70 mV. Maximal glutamate-evoked currents were taken as responses to 100 uM glutamate. 3-HK and NS 3763 were treated for 10 and 3 seconds, respectively, followed by glutamate application.

#### Immune Cell quantification Experiments

For quantification of innate immune cells numbers, *mpeg1:mCherry* animals were outcrossed with *mpx:EGFP^4^* animals to generate double heterozygous transgenic animals. Embryos were injected with either the *pu.1* MO at the single cell stage. At 3 dpf, larvae were anesthetized with tricaine, immobilized in 1% low-melting-point agarose (Sigma) containing tricaine on 35-mm glass-bottom dishes (MatTek), covered with E3 medium containing tricaine and imaged on an epifluorescence microscope (Zeiss Axio Observer). Images were acquired from the whole body of animals and total numbers of GFP^+^ and mCherry^+^ cells per animalwere manually counted using FIJI/image J(*81*).

#### Flow cytometry and fluorescence-activated cell sorter (FACS) analyzes

Homozygous transgenic *Tg(mpeg1:mCherry)* animals were infected with *S.* Typhimurium-GFP, treated or untreated with 3-HK and analyzed by flow cytometry (NovoCyte 3000, Agilent) or fluorescence-activated cell sorter (FACS Aria 3) at 4 hours post infection. Briefly, single cell suspensions were prepared by enzymatic digestion of a group consisting of 5 *mpeg1:mCherry* transgenic animals as previously described(*82*) with one modification including the addition of collagenase 100 mg/ml to the enzymatic digestion. Dead cells were excluded using DAPI (Sigma-Aldrich Inc) or Near-IR Dead Cell Stain (Invitrogen). For bacterial enumeration experiments, sorted cells were collected into 0.3% Triton X-100 in 1X PBS and plated in LB agar plates. Data were analyzed with NovoExpress Version 1.0 (Agilent) and GraphPad Prism 7 (GraphPad).

#### Confocal microscopy imaging of infected zebrafish

Infected embryos were mounted for imaging in LMP agarose (Sigma) containing 0.16 mg/ml tricaine in 35-mm glass-bottom dishes (MatTek), covered with E3 media containing 0.16 mg/ml tricaine and imaged on an A1R (Nikon) confocal microscope using NIS-Elements software (Nikon) as previously described(*83*). All images were analyzed using Fiji (*81*) and were adjusted for brightness and contrast to improve visualization.

#### Transcriptomic analyses

The transcriptome of macrophages isolated from zebrafish embryos were analyzed by RNA-sequencing. Briefly, fluorescently labeled cells were isolated in bulk from a single cell suspension obtained by enzymatic digestion of groups consisting of 5 homozygous Tg(*mpeg1:mCherry*) transgenic animals (8-11 biological replicates/condition) following the protocol as previously described(*82*). cDNA libraries were constructed from a range of 50-800 collected cells with an in-house adaptation of the Smart-seq2 protocol(*84*) in combination with the Nextera XT Library preparation kit (Illumina, Inc., San Diego, CA). Library size and concentration were evaluated using the TapeStation 2200 system (Agilent) and a Qubit fluorometer (Invitrogen) prior to sequencing. Samples were multiplexed and paired-end sequences of 50 bp were generated on a NovaSeq S4 6000 sequencing system (Illumina) generating on average 1.03 x10^8^ reads per sample. Samples were demultiplexed and FASTQ files representing each sample were generated. Any remaining adapter sequences were removed using skewer (0.2.2)(*85*) and FASTQ files were assessed for quality control using FASTQC (0.11.5)(*86*). Reads were aligned to the Ensembl(*87*) zebrafish reference genome GRCz11 using Hisat2 (2.1.0)(*88*) and counts were quantified using HTSeq-Count (0.12.4)(*89*). Differential gene expression analysis was performed using DESeq2 (1.28.1)(*90*) with a significance cut-off of p < 0.05. Gene ontology analyses were performed using gene set enrichment analysis (GSEA)(*91, 92*) and the Kyoto Encyclopedia of Genes and Genomes (KEGG) pathway analyses with a FDR < 0.05 using gseGO, gseKEGG and GSEA, respectively, from ClusterProfiler 4.0(*93, 94*).

#### Tissue penetration of KP metabolites

Tissue penetration of tryptophan was determined by measuring an increase in the downstream tryptophan metabolite, kynurenine (Kyn), by ELISA from zebrafish larval homogenates. Larvae were bathed overnight in 50 uM tryptophan, 100 uM tryptophan, or vehicle (H2O). The following day, larvae were euthanized with tricaine on ice for 10 minutes prior to homogenizing larvae (N=135 per condition) in a final volume of 150 uL ice cold 1X PBS using a motorized Pellet Pestle (Kontes) with disposable polypropylene pellet pestles (Millipore Sigma). Samples were centrifuged at 16,100 x *g* for 5 minutes and the supernatant was filtered through a 40 mm cell filter. Kynurenine concentrations were measured for each sample by ELISA for kynurenine (Abbexa) according to the manufacturer’s instructions. Tissue penetration of 3-HAA, QA, KYNA, and XA were determined by bathing 3 dpf larvae in indicated concentrations of metabolites overnight and measuring the heart rate of each animal per group over a 30 sec interval 19 hours post immersion. Larvae were visualized under brightfield illumination with an inverted microscope (Nikon Eclipse TE2000-S) to more easily visualize the heart ventrally.

#### ROS activity measurements

ROS activity was assayed as previously described(*95*) using the cell-permeant ROS indicator CM-H_2_DCFDA (Life Technologies). Following infection, larvae were arrayed in an optical bottom 96-well plate (Nunc 265301) in wells containing E3, E3 +50 uM 3-HK or 1mM Na-ascorbate as a positive control for antioxidant activity. One hour post infection, fresh CM-H2DCFDA was added to each well at a final concentration of 1ug/ml CM-H2DCFDA and 1% DMSO. The plate was incubated 1 hour in the dark at 28C to facilitate loading of the dye into fish tissues. After 1 hour, the wells containing CM-H2DCFDA were washed once with E3 media. The plate was then incubated in a humidity chamber in the Tecan Spark 10M plate reader (Tecan) overnight with the temperature set at 28C and kinetic reads for fluorescence obtained every hour, with 484/20 nm and 529/20 nm excitation and emission filters, respectively. Wells without fish were used as a negative control for spontaneous oxidation of CM-H2DCFDA.

#### NAD+/ NADH ratio measurements

Larvae (3 dpf) were immersed in either 50 uM 3-HK, 3-HAA, QA or 200 uM β-NMN or vehicle overnight. The following day, larvae were euthanized with tricaine on ice for 5 min. prior to homogenizing four larvae per sample by passage through a 17 gauge needle, as described above for RNA isolation, in a final volume of 100ul 0.2N NaOH and 1% DTAB. The NAD+/NADH ratio from each sample was then assessed using the NAD/NADH-Glo Assay (Promega; Madison, WI) according to the manufacturer’s instructions.

### Cell culture experiments

#### Infection of Primary Murine Bone Marrow Derived Macrophages (BMDMs)

Primary murine BMDMs were isolated from C57/BL6 mice as previously described(*74*) and seeded into 96-well plates at 4.71e4 cells/ well in a total volume of 100ul media (DMEM +20%FBS +5% L-glutamine and 25 ng/ml m-CSF (R&D) and incubated overnight at 37C. The following day, media was removed, and cells were stimulated for 4 hours with *E. coli* LPS (Fluka) at 100ng/ml in complete media described above in a total volume of 100 ul. After 4 hours, the media was removed and replaced with 200ul media (DMEM + 10% FBS) without or with 12.5 uM, 25 uM, or 50 uM 3-HK and mid-log phase bacteria (MOIs 2 and 20) derived from an overnight LB culture of *S.* Typhimurium that had been sub-cultured 1:500 and grown 4 hours with shaking at 37C prior to infection. Plates were centrifuged at 700 x *g* for 15 min at RT to synchronize infections and then incubated at 37C for 30 min. The supernatant was removed from each well and cells were washed twice in 1X PBS before being overlayed with media containing 50ug/ml gentamicin either without or with 12.5 uM, 25 uM, or 50 uM 3-HK and then incubated for a total of 5 hours at 37C before harvesting supernatants to ascertain LDH release using the LDH assay kit (Pierce) according to manufacturer instructions and as previously described(*96*).

#### Infection of Macrophage/ Monocyte Cell Lines

For infection of cell lines (RAW264, J774 or U937 cells (ATCC)), cells were seeded in 24 well plates (1e5 cells/ well, RAW264 and U937; 5e5 cells/well, J774) in a total volume of 0.5ml media (DMEM + 10% FBS, RAW264 and J774; RPMI with 10% FBS, U937) per well for bacterial burden determinations. For host cell survival determinations, cells (5e4 cells/ well, J774 and U937; 3.2e4 cells/well, RAW264) were seeded into 96-well plates in a total volume of 100uL. For infections with U937 cells, a final concentration of 10nM PMA was added to media when cells were seeded into assay plates to facilitate differentiation and adhesion of these cells to the plate bottom. Seeded plates were incubated overnight at 37C and the following day, cells were infected with *S.* Typhimurium SL1344 derived from a stationary phase culture grown overnight in LB or sub-cultured and grown to log phase as described above. For infections with stationary phase bacteria, bacteria were opsonized 1:1 with normal mouse serum (RAW264 and J774, Jackson ImmunoResearch) or normal human serum (U937,) at 37C for 15 min. prior to infection. Infections with stationary and log phase bacteria were carried out as described above for BMDMs. Following the addition of bacteria to cells, plates were centrifuged at 165 x g for 5 min. to synchronize infections and plates were incubated for 30 min. at 37C to allow uptake of bacteria. After 30 min., the supernatant was removed from each well and cells were washed twice in 1X PBS before being overlayed with media with gentamicin (50-100ug/ml) either without or with 12.5 uM, 25 uM, or 50 uM 3-HK. Plates were incubated a further 90 min. after which the media was again removed from each well and replaced with media containing a lower concentration of gentamicin (5-20ug/ml) either without or with 12.5 uM, 25 uM, or 50 uM 3-HK. For bacterial burden measurements, cells were lysed 24 HPI to enumerate colonizing bacteria. Briefly, media was removed from each well and the well overlayed with 1ml 1XPBS + 0.1% Triton X-100. Each well was pipetted up and down 20X to lyse mammalian cells and the lysate collected, vortexed heavily, diluted 1:1000 in PBS before to plating to LB-strep (25ug/ml) plates for CFU determination measurements. Assays for host cell viability following infection were carried out with Cell Titer Glo (Promega; J774 and U937) or the LDH assay kit (Pierce; RAW264) according to the manufacturer’s instructions and as previously described(*96*). RAW264 MOIs of 10 (LDH and burden assays) and 20 (LDH assay only). J774 MOIs:

U937 MOI:

#### Construction of P. aeruginosa PA14ΔoprM

An in-frame clean deletion of the *oprM* gene was created in strain background *P. aeruginosa* PA14 (gift from F. Ausubel, MGH) using standard allelic exchange techniques. Briefly, the flanking regions 977 bp upstream and 922 bp downstream of the *oprM* gene were amplified with primers AC540 and AC528 and AC541 and AC531 and the left and right flanking region amplicons, which included the start and stop codons for *oprM*, were stitched together using overlap extension PCR using primers AC528 and AC531 that had been phosphorylated with T4 Polynucleotide kinase (NEB). The resulting PCR product was then blunt-cloned into pEX18Ap(*97*), which had been linearized with EcoRI and HindIII, blunt end-polished with T4 DNA polymerase (NEB) and dephosphorylated with Antarctic phosphatase (NEB). The resulting plasmid, pEX18Amp_oprM2 was introduced into *E. coli* SM10 by electroporation and mated with *P. aeruginosa* PA14. Single-crossover merodiploid exconjugants were selected based on carbenicillin (150ug/ml) and irgasan (15ug/ml) resistance and double cross-over recombinants were isolated by plating the merodiploids to LB agar with 6% sucrose to force the removal of vector DNA containing the *sacB* gene. In-frame deletion of *oprM* was confirmed by PCR amplification of the flanking regions of the target gene with primers AC554 and AC555 as well as AC556 and AC557 and sequencing the resulting products.

#### Primer sequences

PA14*1′oprM strain construction*

AC540: 5’- CCAAATGCCAAATCGTGCCATATCATTGCCCCTTTTCGACGGA-3’

AC528: 5’- TTGAATTCAGTACAAGCTGGAGATCGACGAC-3’

AC541: 5’- GCACGATTTGGCATTTGGTGATCGCCTTCCGCGCCATGCAAGAA-3’

AC531: 5’- ATGGATCCAGAAGATGACCAGACAGCGTATTC-3’

AC554: 5’- GGGTACCGCTGTTCTACGTG-3’

AC555: 5’- AAGACTACGTCCACCGCATC-3’

AC556: 5’- GCAACAAGTTCCTCATGCTC-3’

AC557: 5’-AGCCGGTGGTGATCAACAAG-3’

*grik1b*, *grik1a*, g*rik4* and *grik5* transcript cloning for electrophysiology experiments

SC42: 5’- ATGAGTGACTCCCATGAGCACC-3’

AC936: 5’- ATGAGCACCAGTGCGAGTGA-3’

SC8: 5’- ATGAAGTTTTGGCTTTTAACCTCTTTTGATAGC-3’

AC935: 5’-CTAACGCGAAGAACATATGTC-3’

CC189: 5’- TTGCAGGATCCCATCGATTCATGCCGGAACTGCCAGCG-3’

CC190: 5’- CACTATAGGGCTGCAGAATCTCATTTTTGCTCCCCCATTAGGTC-3’

SC179: 5’- GCCAACTTTGTACAAAAAAGCAGGCTCCGAATTCGCCCTTATGCTCGCAGCCTGGGC-3’

SC180: 5’- GCCAACTTTGTACAAGAAAGCTGGGTCGAATTTTATTCGTGTTCACTGTTGTTGGTGC C-3’

#### Statistical analysis

Statistical analyses were carried out using GraphPad Prism 7 (GraphPad).

## REFERENCES

1. D. Bumann, Heterogeneous host-pathogen encounters: act locally, think globally. Cell host & microbe 17, 13–19 (2015).

2. R. Avraham, Untangling Cellular Host-Pathogen Encounters at Infection Bottlenecks. Infection and immunity, e0043822 (2023).

3. S. Masud, V. Torraca, A. H. Meijer, Modeling Infectious Diseases in the Context of a Developing Immune System. Curr Top Dev Biol 124, 277–329 (2017).

4. I. Cervenka, L. Z. Agudelo, J. L. Ruas, Kynurenines: Tryptophan’s metabolites in exercise, inflammation, and mental health. Science 357, (2017).

5. Y. S. Huang, J. Ogbechi, F. I. Clanchy, R. O. Williams, T. W. Stone, IDO and Kynurenine Metabolites in Peripheral and CNS Disorders. Front Immunol 11, 388 (2020).

6. B. Ricciuti, G. C. Leonardi, P. Puccetti, F. Fallarino, V. Bianconi, A. Sahebkar, S. Baglivo, R. Chiari, M. Pirro, Targeting indoleamine-2,3-dioxygenase in cancer: Scientific rationale and clinical evidence. Pharmacol Ther 196, 105–116 (2019).

7. M. Modoux, N. Rolhion, S. Mani, H. Sokol, Tryptophan Metabolism as a Pharmacological Target. Trends Pharmacol Sci 42, 60–73 (2021).

8. M. Platten, E. A. A. Nollen, U. F. Rohrig, F. Fallarino, C. A. Opitz, Tryptophan metabolism as a common therapeutic target in cancer, neurodegeneration and beyond. Nat Rev Drug Discov 18, 379–401 (2019).

9. A. E. Clatworthy, J. S. Lee, M. Leibman, Z. Kostun, A. J. Davidson, D. T. Hung, Pseudomonas aeruginosa infection of zebrafish involves both host and pathogen determinants. Infection and immunity 77, 1293–1303 (2009).

10. P. I. Fields, R. V. Swanson, C. G. Haidaris, F. Heffron, Mutants of Salmonella typhimurium that cannot survive within the macrophage are avirulent. Proceedings of the National Academy of Sciences of the United States of America 83, 5189–5193 (1986).

11. K. Y. Leung, B. B. Finlay, Intracellular replication is essential for the virulence of Salmonella typhimurium. Proceedings of the National Academy of Sciences of the United States of America 88, 11470–11474 (1991).

12. A. M. van der Sar, R. J. Musters, F. J. van Eeden, B. J. Appelmelk, C. M. Vandenbroucke-Grauls, W. Bitter, Zebrafish embryos as a model host for the real time analysis of Salmonella typhimurium infections. Cell Microbiol 5, 601–611 (2003).

13. J. Rhodes, A. Hagen, K. Hsu, M. Deng, T. X. Liu, A. T. Look, J. P. Kanki, Interplay of pu.1 and gata1 determines myelo-erythroid progenitor cell fate in zebrafish. Dev Cell 8, 97–108 (2005).

14. T. K. Prajsnar, V. T. Cunliffe, S. J. Foster, S. A. Renshaw, A novel vertebrate model of Staphylococcus aureus infection reveals phagocyte-dependent resistance of zebrafish to non-host specialized pathogens. Cell Microbiol 10, 2312–2325 (2008).

15. S. Masud, T. K. Prajsnar, V. Torraca, G. E. M. Lamers, M. Benning, M. Van Der Vaart, A. H. Meijer, Macrophages target Salmonella by Lc3-associated phagocytosis in a systemic infection model. Autophagy 15, 796–812 (2019).

16. J. M. Farrow, 3rd, E. C. Pesci, Two distinct pathways supply anthranilate as a precursor of the Pseudomonas quinolone signal. Journal of bacteriology 189, 3425–3433 (2007).

17. O. Kurnasov, V. Goral, K. Colabroy, S. Gerdes, S. Anantha, A. Osterman, T. P. Begley, NAD biosynthesis: identification of the tryptophan to quinolinate pathway in bacteria. Chem Biol 10, 1195–1204 (2003).

18. X. D. Wang, F. M. Notarangelo, J. Z. Wang, R. Schwarcz, Kynurenic acid and 3-hydroxykynurenine production from D-kynurenine in mice. Brain Res 1455, 1–9 (2012).

19. S. Kumar, D. Jaller, B. Patel, J. M. LaLonde, J. B. DuHadaway, W. P. Malachowski, G. C. Prendergast, A. J. Muller, Structure based development of phenylimidazole-derived inhibitors of indoleamine 2,3-dioxygenase. J Med Chem 51, 4968–4977 (2008).

20. S. Rover, A. M. Cesura, P. Huguenin, R. Kettler, A. Szente, Synthesis and biochemical evaluation of N-(4-phenylthiazol-2-yl)benzenesulfonamides as high-affinity inhibitors of kynurenine 3-hydroxylase. J Med Chem 40, 4378–4385 (1997).

21. T. W. Stone, R. O. Williams, Modulation of T cells by tryptophan metabolites in the kynurenine pathway. Trends Pharmacol Sci 44, 442–456 (2023).

22. L. E. Goldstein, M. C. Leopold, X. Huang, C. S. Atwood, A. J. Saunders, M. Hartshorn, J. T. Lim, K. Y. Faget, J. A. Muffat, R. C. Scarpa, L. T. Chylack, Jr., E. F. Bowden, R. E. Tanzi, A. I. Bush, 3-Hydroxykynurenine and 3-hydroxyanthranilic acid generate hydrogen peroxide and promote alpha-crystallin cross-linking by metal ion reduction. Biochemistry 39, 7266–7275 (2000).

23. S. Okuda, N. Nishiyama, H. Saito, H. Katsuki, Hydrogen peroxide-mediated neuronal cell death induced by an endogenous neurotoxin, 3-hydroxykynurenine. Proceedings of the National Academy of Sciences of the United States of America 93, 12553–12558 (1996).

24. S. Okuda, N. Nishiyama, H. Saito, H. Katsuki, 3-Hydroxykynurenine, an endogenous oxidative stress generator, causes neuronal cell death with apoptotic features and region selectivity. J Neurochem 70, 299–307 (1998).

25. S. Christen, E. Peterhans, R. Stocker, Antioxidant activities of some tryptophan metabolites: possible implication for inflammatory diseases. Proceedings of the National Academy of Sciences of the United States of America 87, 2506–2510 (1990).

26. A. L. Colin-Gonzalez, P. D. Maldonado, A. Santamaria, 3-Hydroxykynurenine: an intriguing molecule exerting dual actions in the central nervous system. Neurotoxicology 34, 189–204 (2013).

27. G. McClarty, H. D. Caldwell, D. E. Nelson, Chlamydial interferon gamma immune evasion influences infection tropism. Curr Opin Microbiol 10, 47–51 (2007).

28. E. R. Pfefferkorn, Interferon gamma blocks the growth of Toxoplasma gondii in human fibroblasts by inducing the host cells to degrade tryptophan. Proceedings of the National Academy of Sciences of the United States of America 81, 908–912 (1984).

29. L. D. Sibley, M. Messina, I. R. Niesman, Stable DNA transformation in the obligate intracellular parasite Toxoplasma gondii by complementation of tryptophan auxotrophy. Proceedings of the National Academy of Sciences of the United States of America 91, 5508–5512 (1994).

30. K. Peng, D. M. Monack, Indoleamine 2,3-dioxygenase 1 is a lung-specific innate immune defense mechanism that inhibits growth of Francisella tularensis tryptophan auxotrophs. Infection and immunity 78, 2723–2733 (2010).

31. Q. Li, J. L. Harden, C. D. Anderson, N. K. Egilmez, Tolerogenic Phenotype of IFN-gamma-Induced IDO+ Dendritic Cells Is Maintained via an Autocrine IDO-Kynurenine/AhR-IDO Loop. J Immunol 197, 962–970 (2016).

32. J. D. Mezrich, J. H. Fechner, X. Zhang, B. P. Johnson, W. J. Burlingham, C. A. Bradfield, An interaction between kynurenine and the aryl hydrocarbon receptor can generate regulatory T cells. J Immunol 185, 3190–3198 (2010).

33. A. Puyskens, A. Stinn, M. van der Vaart, A. Kreuchwig, J. Protze, G. Pei, M. Klemm, U. Guhlich-Bornhof, R. Hurwitz, G. Krishnamoorthy, M. Schaaf, G. Krause, A. H. Meijer, S. H. E. Kaufmann, P. Moura-Alves, Aryl Hydrocarbon Receptor Modulation by Tuberculosis Drugs Impairs Host Defense and Treatment Outcomes. Cell host & microbe 27, 238–248 e237 (2020).

34. V. M. D’Costa, V. Braun, M. Landekic, R. Shi, A. Proteau, L. McDonald, M. Cygler, S. Grinstein, J. H. Brumell, Salmonella Disrupts Host Endocytic Trafficking by SopD2-Mediated Inhibition of Rab7. Cell Rep 12, 1508–1518 (2015).

35. M. Doerflinger, Y. Deng, P. Whitney, R. Salvamoser, S. Engel, A. J. Kueh, L. Tai, A. Bachem, E. Gressier, N. D. Geoghegan, S. Wilcox, K. L. Rogers, A. L. Garnham, M. A. Dengler, S. M. Bader, G. Ebert, J. S. Pearson, D. De Nardo, N. Wang, C. Yang, M. Pereira, C. E. Bryant, R. A. Strugnell, J. E. Vince, M. Pellegrini, A. Strasser, S. Bedoui, M. J. Herold, Flexible Usage and Interconnectivity of Diverse Cell Death Pathways Protect against Intracellular Infection. Immunity 53, 533–547 e537 (2020).

36. D. G. McEwan, B. Richter, B. Claudi, C. Wigge, P. Wild, H. Farhan, K. McGourty, F. P. Coxon, M. Franz-Wachtel, B. Perdu, M. Akutsu, A. Habermann, A. Kirchof, M. H. Helfrich, P. R. Odgren, W. Van Hul, A. S. Frangakis, K. Rajalingam, B. Macek, D. W. Holden, D. Bumann, I. Dikic, PLEKHM1 regulates Salmonella-containing vacuole biogenesis and infection. Cell host & microbe 17, 58–71 (2015).

37. S. L. Petersen, T. T. Chen, D. A. Lawrence, S. A. Marsters, F. Gonzalvez, A. Ashkenazi, TRAF2 is a biologically important necroptosis suppressor. Cell Death Differ 22, 1846–1857 (2015).

38. I. Dikic, Z. Elazar, Mechanism and medical implications of mammalian autophagy. Nat Rev Mol Cell Biol 19, 349–364 (2018).

39. J. E. Casanova, Bacterial Autophagy: Offense and Defense at the Host-Pathogen Interface. Cell Mol Gastroenterol Hepatol 4, 237–243 (2017).

40. S. Masud, L. van der Burg, L. Storm, T. K. Prajsnar, A. H. Meijer, Rubicon-Dependent Lc3 Recruitment to Salmonella-Containing Phagosomes Is a Host Defense Mechanism Triggered Independently From Major Bacterial Virulence Factors. Front Cell Infect Microbiol 9, 279 (2019).

41. C. P. Knubel, F. F. Martinez, R. E. Fretes, C. Diaz Lujan, M. G. Theumer, L. Cervi, C. C. Motran, Indoleamine 2,3-dioxigenase (IDO) is critical for host resistance against Trypanosoma cruzi. FASEB J 24, 2689–2701 (2010).

42. C. R. MacKenzie, K. Heseler, A. Muller, W. Daubener, Role of indoleamine 2,3-dioxygenase in antimicrobial defence and immuno-regulation: tryptophan depletion versus production of toxic kynurenines. Curr Drug Metab 8, 237–244 (2007).

43. A. Nino-Castro, Z. Abdullah, A. Popov, Y. Thabet, M. Beyer, P. Knolle, E. Domann, T. Chakraborty, S. V. Schmidt, J. L. Schultze, The IDO1-induced kynurenines play a major role in the antimicrobial effect of human myeloid cells against Listeria monocytogenes. Innate Immun 20, 401–411 (2014).

44. C. J. Blohmke, T. C. Darton, C. Jones, N. M. Suarez, C. S. Waddington, B. Angus, L. Zhou, J. Hill, S. Clare, L. Kane, S. Mukhopadhyay, F. Schreiber, M. A. Duque-Correa, J. C. Wright, T. I. Roumeliotis, L. Yu, J. S. Choudhary, A. Mejias, O. Ramilo, M. Shanyinde, M. B. Sztein, R. A. Kingsley, S. Lockhart, M. M. Levine, D. J. Lynn, G. Dougan, A. J. Pollard, Interferon-driven alterations of the host’s amino acid metabolism in the pathogenesis of typhoid fever. J Exp Med 213, 1061–1077 (2016).

45. V. R. Clarke, S. M. Molchanova, T. Hirvonen, T. Taira, S. E. Lauri, Activity-dependent upregulation of presynaptic kainate receptors at immature CA3-CA1 synapses. J Neurosci 34, 16902–16916 (2014).

46. P. Follesa, M. K. Ticku, NMDA receptor upregulation: molecular studies in cultured mouse cortical neurons after chronic antagonist exposure. J Neurosci 16, 2172–2178 (1996).

47. M. J. Keiser, B. L. Roth, B. N. Armbruster, P. Ernsberger, J. J. Irwin, B. K. Shoichet, Relating protein pharmacology by ligand chemistry. Nat Biotechnol 25, 197–206 (2007).

48. A. K. Bhandage, Z. Jin, C. Hellgren, S. V. Korol, K. Nowak, L. Williamsson, I. Sundstrom-Poromaa, B. Birnir, AMPA, NMDA and kainate glutamate receptor subunits are expressed in human peripheral blood mononuclear cells (PBMCs) where the expression of GluK4 is altered by pregnancy and GluN2D by depression in pregnant women. J Neuroimmunol 305, 51–58 (2017).

49. N. Inagaki, H. Kuromi, T. Gonoi, Y. Okamoto, H. Ishida, Y. Seino, T. Kaneko, T. Iwanaga, S. Seino, Expression and role of ionotropic glutamate receptors in pancreatic islet cells. FASEB J 9, 686–691 (1995).

50. G. Lombardi, C. Dianzani, G. Miglio, P. L. Canonico, R. Fantozzi, Characterization of ionotropic glutamate receptors in human lymphocytes. Br J Pharmacol 133, 936–944 (2001).

51. H. Sun, A. Swaim, J. E. Herrera, D. Becker, L. Becker, K. Srivastava, L. E. Thompson, M. R. Shero, A. Perez-Tamayo, B. Suktitipat, R. Mathias, A. Contractor, N. Faraday, C. N. Morrell, Platelet kainate receptor signaling promotes thrombosis by stimulating cyclooxygenase activation. Circ Res 105, 595–603 (2009).

52. J. Lerma, J. M. Marques, Kainate receptors in health and disease. Neuron 80, 292–311 (2013).

53. S. Valbuena, J. Lerma, Non-canonical Signaling, the Hidden Life of Ligand-Gated Ion Channels. Neuron 92, 316–329 (2016).

54. S. Tomita, P. E. Castillo, Neto1 and Neto2: auxiliary subunits that determine key properties of native kainate receptors. J Physiol 590, 2217–2223 (2012).

55. I. H. Greger, J. F. Watson, S. G. Cull-Candy, Structural and Functional Architecture of AMPA-Type Glutamate Receptors and Their Auxiliary Proteins. Neuron 94, 713–730 (2017).

56. A. C. Jackson, R. A. Nicoll, The expanding social network of ionotropic glutamate receptors: TARPs and other transmembrane auxiliary subunits. Neuron 70, 178–199 (2011).

57. J. K. Christensen, T. Varming, P. K. Ahring, T. D. Jorgensen, E. O. Nielsen, In vitro characterization of 5-carboxyl-2,4-di-benzamidobenzoic acid (NS3763), a noncompetitive antagonist of GLUK5 receptors. J Pharmacol Exp Ther 309, 1003–1010 (2004).

58. T. Honore, S. N. Davies, J. Drejer, E. J. Fletcher, P. Jacobsen, D. Lodge, F. E. Nielsen, Quinoxalinediones: potent competitive non-NMDA glutamate receptor antagonists. Science 241, 701–703 (1988).

59. J. Valgeirsson, E. O. Nielsen, D. Peters, T. Varming, C. Mathiesen, A. S. Kristensen, U. Madsen, 2-arylureidobenzoic acids: selective noncompetitive antagonists for the homomeric kainate receptor subtype GluR5. J Med Chem 46, 5834–5843 (2003).

60. J. K. Christensen, A. V. Paternain, S. Selak, P. K. Ahring, J. Lerma, A mosaic of functional kainate receptors in hippocampal interneurons. J Neurosci 24, 8986–8993 (2004).

61. D. Aggad, C. Stein, D. Sieger, M. Mazel, P. Boudinot, P. Herbomel, J. P. Levraud, G. Lutfalla, M. Leptin, In vivo analysis of Ifn-gamma1 and Ifn-gamma2 signaling in zebrafish. J Immunol 185, 6774–6782 (2010).

62. L. Jiang, P. Wang, X. Song, H. Zhang, S. Ma, J. Wang, W. Li, R. Lv, X. Liu, S. Ma, J. Yan, H. Zhou, D. Huang, Z. Cheng, C. Yang, L. Feng, L. Wang, Salmonella Typhimurium reprograms macrophage metabolism via T3SS effector SopE2 to promote intracellular replication and virulence. Nat Commun 12, 879 (2021).

63. G. Rosenberg, D. Yehezkel, D. Hoffman, C. C. Mattioli, M. Fremder, H. Ben-Arosh, L. Vainman, N. Nissani, S. Hen-Avivi, S. Brenner, M. Itkin, S. Malitsky, E. Ohana, N. B. Ben-Moshe, R. Avraham, Host succinate is an activation signal for Salmonella virulence during intracellular infection. Science 371, 400–405 (2021).

64. D. E. Nelson, D. P. Virok, H. Wood, C. Roshick, R. M. Johnson, W. M. Whitmire, D. D. Crane, O. Steele-Mortimer, L. Kari, G. McClarty, H. D. Caldwell, Chlamydial IFN-gamma immune evasion is linked to host infection tropism. Proceedings of the National Academy of Sciences of the United States of America 102, 10658–10663 (2005).

65. C. P. Knubel, C. Insfran, F. F. Martinez, C. Diaz Lujan, R. E. Fretes, M. G. Theumer, L. Cervi, C. C. Motran, 3-Hydroxykynurenine, a Tryptophan Metabolite Generated during the Infection, Is Active Against Trypanosoma cruzi. ACS Med Chem Lett 8, 757–761 (2017).

66. P. Guidetti, R. Schwarcz, 3-Hydroxykynurenine potentiates quinolinate but not NMDA toxicity in the rat striatum. Eur J Neurosci 11, 3857–3863 (1999).

67. M. N. Perkins, T. W. Stone, An iontophoretic investigation of the actions of convulsant kynurenines and their interaction with the endogenous excitant quinolinic acid. Brain Res 247, 184–187 (1982).

68. K. S. Elmslie, D. Yoshikami, Effects of kynurenate on root potentials evoked by synaptic activity and amino acids in the frog spinal cord. Brain Res 330, 265–272 (1985).

69. M. Y. Min, Z. Melyan, D. M. Kullmann, Synaptically released glutamate reduces gamma-aminobutyric acid (GABA)ergic inhibition in the hippocampus via kainate receptors. Proceedings of the National Academy of Sciences of the United States of America 96, 9932–9937 (1999).

70. C. Prescott, A. M. Weeks, K. J. Staley, K. M. Partin, Kynurenic acid has a dual action on AMPA receptor responses. Neurosci Lett 402, 108–112 (2006).

71. A. Contractor, C. Mulle, G. T. Swanson, Kainate receptors coming of age: milestones of two decades of research. Trends Neurosci 34, 154–163 (2011).

72. I. D. Jung, M. G. Lee, J. H. Chang, J. S. Lee, Y. I. Jeong, C. M. Lee, W. S. Park, J. Han, S. K. Seo, S. Y. Lee, Y. M. Park, Blockade of indoleamine 2,3-dioxygenase protects mice against lipopolysaccharide-induced endotoxin shock. J Immunol 182, 3146–3154 (2009).

73. C. Skouras, X. Zheng, M. Binnie, N. Z. Homer, T. B. Murray, D. Robertson, L. Briody, F. Paterson, H. Spence, L. Derr, A. J. Hayes, A. Tsoumanis, D. Lyster, R. W. Parks, O. J. Garden, J. P. Iredale, I. J. Uings, J. Liddle, W. L. Wright, G. Dukes, S. P. Webster, D. J. Mole, Increased levels of 3-hydroxykynurenine parallel disease severity in human acute pancreatitis. Sci Rep 6, 33951 (2016).

74. R. Avraham, N. Haseley, D. Brown, C. Penaranda, H. B. Jijon, J. J. Trombetta, R. Satija, A. K. Shalek, R. J. Xavier, A. Regev, D. T. Hung, Pathogen Cell-to-Cell Variability Drives Heterogeneity in Host Immune Responses. Cell 162, 1309–1321 (2015).

75. S. A. Stanley, T. Kawate, N. Iwase, M. Shimizu, A. E. Clatworthy, E. Kazyanskaya, J. C. Sacchettini, T. R. Ioerger, N. A. Siddiqi, S. Minami, J. A. Aquadro, S. S. Grant, E. J. Rubin, D. T. Hung, Diarylcoumarins inhibit mycolic acid biosynthesis and kill Mycobacterium tuberculosis by targeting FadD32. Proceedings of the National Academy of Sciences of the United States of America 110, 11565–11570 (2013).

76. S. Wellington, P. P. Nag, K. Michalska, S. E. Johnston, R. P. Jedrzejczak, V. K. Kaushik, A. E. Clatworthy, N. Siddiqi, P. McCarren, B. Bajrami, N. I. Maltseva, S. Combs, S. L. Fisher, A. Joachimiak, S. L. Schreiber, D. T. Hung, A small-molecule allosteric inhibitor of Mycobacterium tuberculosis tryptophan synthase. Nature chemical biology 13, 943–950 (2017).

77. F. Ellett, L. Pase, J. W. Hayman, A. Andrianopoulos, G. J. Lieschke, mpeg1 promoter transgenes direct macrophage-lineage expression in zebrafish. Blood 117, e49–56 (2011).

78. S. A. Renshaw, C. A. Loynes, D. M. Trushell, S. Elworthy, P. W. Ingham, M. K. Whyte, A transgenic zebrafish model of neutrophilic inflammation. Blood 108, 3976–3978 (2006).

79. N. Van Rooijen, A. Sanders, Liposome mediated depletion of macrophages: mechanism of action, preparation of liposomes and applications. J Immunol Methods 174, 83–93 (1994).

80. W. Zhang, F. St-Gelais, C. P. Grabner, J. C. Trinidad, A. Sumioka, M. Morimoto-Tomita, K. S. Kim, C. Straub, A. L. Burlingame, J. R. Howe, S. Tomita, A transmembrane accessory subunit that modulates kainate-type glutamate receptors. Neuron 61, 385–396 (2009).

81. J. Schindelin, I. Arganda-Carreras, E. Frise, V. Kaynig, M. Longair, T. Pietzsch, S. Preibisch, C. Rueden, S. Saalfeld, B. Schmid, J. Y. Tinevez, D. J. White, V. Hartenstein, K. Eliceiri, P. Tomancak, A. Cardona, Fiji: an open-source platform for biological-image analysis. Nature methods 9, 676–682 (2012).

82. V. E. Gallardo, J. Liang, M. Behra, A. Elkahloun, E. J. Villablanca, V. Russo, M. L. Allende, S. M. Burgess, Molecular dissection of the migrating posterior lateral line primordium during early development in zebrafish. BMC Dev Biol 10, 120 (2010).

83. O. Renaud, P. Herbomel, K. Kissa, Studying cell behavior in whole zebrafish embryos by confocal live imaging: application to hematopoietic stem cells. Nature protocols 6, 1897–1904 (2011).

84. S. Picelli, O. R. Faridani, A. K. Bjorklund, G. Winberg, S. Sagasser, R. Sandberg, Full-length RNA-seq from single cells using Smart-seq2. Nature protocols 9, 171–181 (2014).

85. H. Jiang, R. Lei, S. W. Ding, S. Zhu, Skewer: a fast and accurate adapter trimmer for next-generation sequencing paired-end reads. BMC Bioinformatics 15, 182 (2014).

86 .(2015).

87 . K. L. Howe, P. Achuthan, J. Allen, J. Allen, J. Alvarez-Jarreta, M. R. Amode, I. M. Armean, A. G. Azov, R. Bennett, J. Bhai, K. Billis, S. Boddu, M. Charkhchi, C. Cummins, L. Da Rin Fioretto, C. Davidson, K. Dodiya, B. El Houdaigui, R. Fatima, A. Gall, C. Garcia Giron, T. Grego, C. Guijarro-Clarke, L. Haggerty, A. Hemrom, T. Hourlier, O. G. Izuogu, T. Juettemann, V. Kaikala, M. Kay, I. Lavidas, T. Le, D. Lemos, J. Gonzalez Martinez, J. C. Marugan, T. Maurel, A. C. McMahon, S. Mohanan, B. Moore, M. Muffato, D. N. Oheh, D. Paraschas, A. Parker, A. Parton, I. Prosovetskaia, M. P. Sakthivel, A. I. A. Salam, B. M. Schmitt, H. Schuilenburg, D. Sheppard, E. Steed, M. Szpak, M. Szuba, K. Taylor, A. Thormann, G. Threadgold, B. Walts, A. Winterbottom, M. Chakiachvili, A. Chaubal, N. De Silva, B. Flint, A. Frankish, S. E. Hunt, I. I., Grn. Langridge, J. E. Loveland, F. J. Martin, J. M. Mudge, J. Morales, E. Perry, M. Ruffier, J. Tate, D. Thybert, S. J. Trevanion, F. Cunningham, A. D. Yates, D. R. Zerbino, P. Flicek, Ensembl 2021. Nucleic acids research 49, D884–D891 (2021).

88. D. Kim, J. M. Paggi, C. Park, C. Bennett, S. L. Salzberg, Graph-based genome alignment and genotyping with HISAT2 and HISAT-genotype. Nat Biotechnol 37, 907–915 (2019).

89. S. Anders, P. T. Pyl, W. Huber, HTSeq--a Python framework to work with high-throughput sequencing data. Bioinformatics 31, 166–169 (2015).

90. M. I. Love, W. Huber, S. Anders, Moderated estimation of fold change and dispersion for RNA-seq data with DESeq2. Genome biology 15, 550 (2014).

91. V. K. Mootha, C. M. Lindgren, K. F. Eriksson, A. Subramanian, S. Sihag, J. Lehar, P. Puigserver, E. Carlsson, M. Ridderstrale, E. Laurila, N. Houstis, M. J. Daly, N. Patterson, J. P. Mesirov, T. R. Golub, P. Tamayo, B. Spiegelman, E. S. Lander, J. N. Hirschhorn, D. Altshuler, L. C. Groop, PGC-1alpha-responsive genes involved in oxidative phosphorylation are coordinately downregulated in human diabetes. Nat Genet 34, 267–273 (2003).

92. A. Subramanian, P. Tamayo, V. K. Mootha, S. Mukherjee, B. L. Ebert, M. A. Gillette, A. Paulovich, S. L. Pomeroy, T. R. Golub, E. S. Lander, J. P. Mesirov, Gene set enrichment analysis: a knowledge-based approach for interpreting genome-wide expression profiles. Proceedings of the National Academy of Sciences of the United States of America 102, 15545–15550 (2005).

93. T. Wu, E. Hu, S. Xu, M. Chen, P. Guo, Z. Dai, T. Feng, L. Zhou, W. Tang, L. Zhan, X. Fu, S. Liu, X. Bo, G. Yu, clusterProfiler 4.0: A universal enrichment tool for interpreting omics data. Innovation (Camb) 2, 100141 (2021).

94. G. Yu, L. G. Wang, Y. Han, Q. Y. He, clusterProfiler: an R package for comparing biological themes among gene clusters. OMICS 16, 284–287 (2012).

95. F. J. Roca, L. Ramakrishnan, TNF dually mediates resistance and susceptibility to mycobacteria via mitochondrial reactive oxygen species. Cell 153, 521–534 (2013).

96. P. Broz, D. M. Monack, Measuring inflammasome activation in response to bacterial infection. Methods in molecular biology 1040, 65–84 (2013).

97. T. T. Hoang, R. R. Karkhoff-Schweizer, A. J. Kutchma, H. P. Schweizer, A broad-host-range Flp-FRT recombination system for site-specific excision of chromosomally-located DNA sequences: application for isolation of unmarked Pseudomonas aeruginosa mutants. Gene 212, 77–86 (1998).

