## Supplemental Information for "A tryptophan metabolite modulates the host response to bacterial infection via kainate receptors"

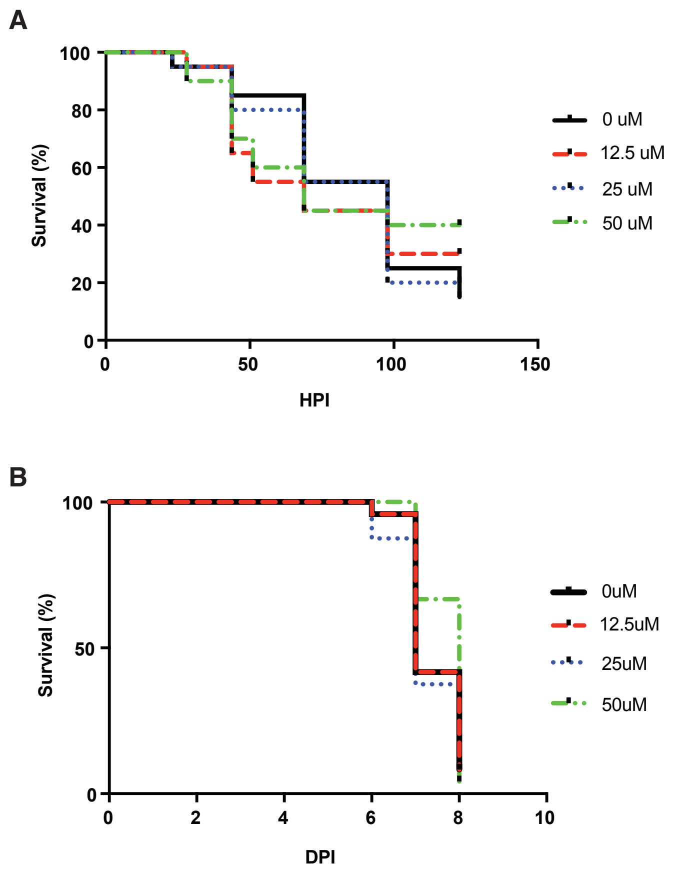


**fig. S1. 3-HK does not promote survival following infection with *S. aureus* or *M. marinum*.** Embryos (48 hpf) were infected intravenously with either *S. aureus* (**A;** 7017 +/- 1754 CFU) or *M. marinum* (**B;** 59 +/- 6 CFU), treated with indicated concentrations of 3-HK and monitored over time for survival. (N=20 embryos/ condition (**A**) or N=24 embryos/ condition (**B**); data are representative of 3 independent experiments). hpi, hours post infection; dpi, days post infection.

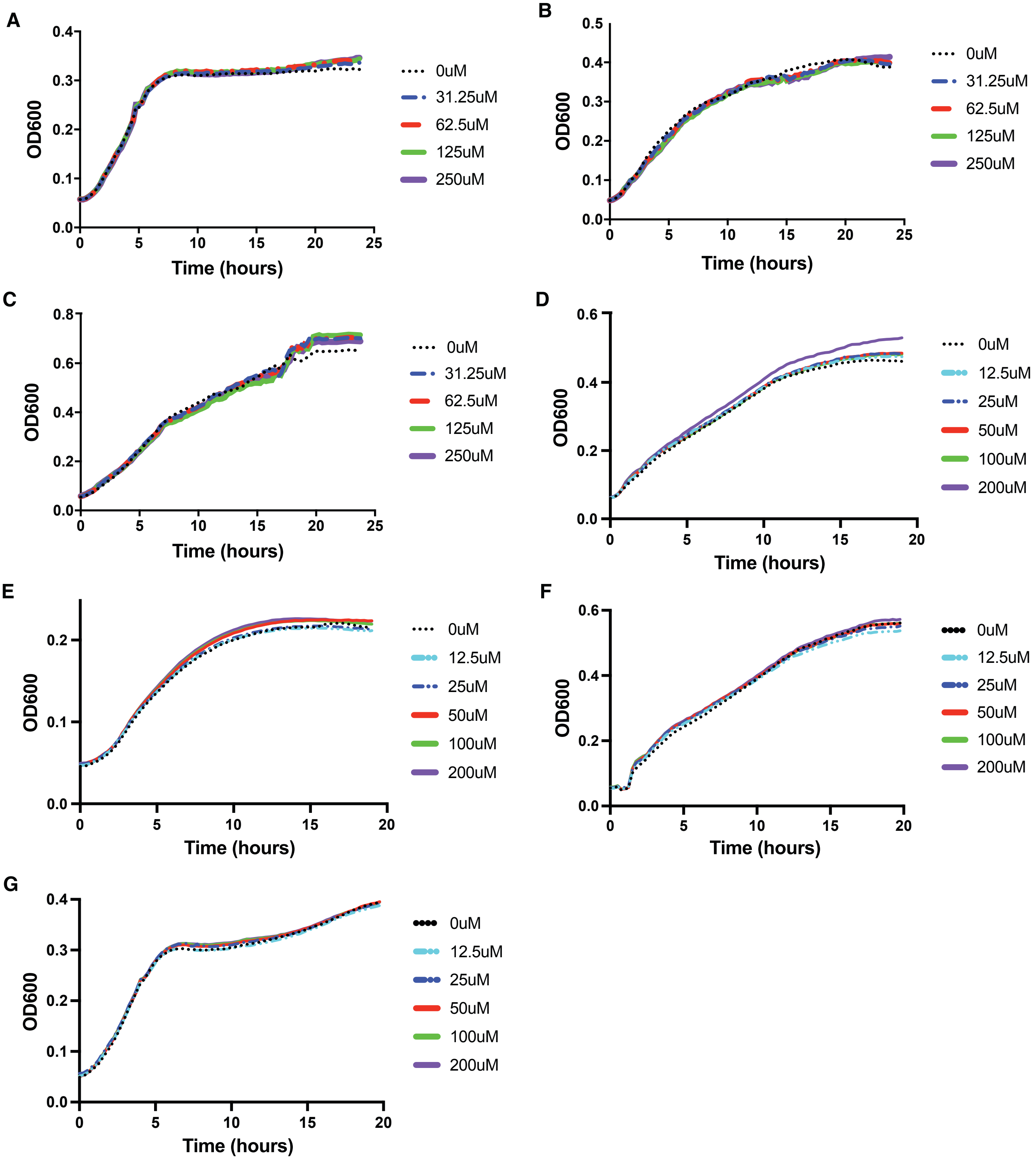


**fig. S2. 3-HK, NS 3763 and CNQX do not alter bacterial growth *in vitro*.** Bacterial strains (*S*. Typhimurium SL1344 (**A**,**D**,**E**,**F**,**G**) and *P. aeruginosa* PA14Δ*oprM* (**B**,**C**) were grown in M9 (**A, B, E, G**) or LB (**C**,**D**,**F**) in indicated concentrations of 3-HK (**A**-**C**), NS 3763 (**D**,**E**) or CNQX (**F**,**G**).

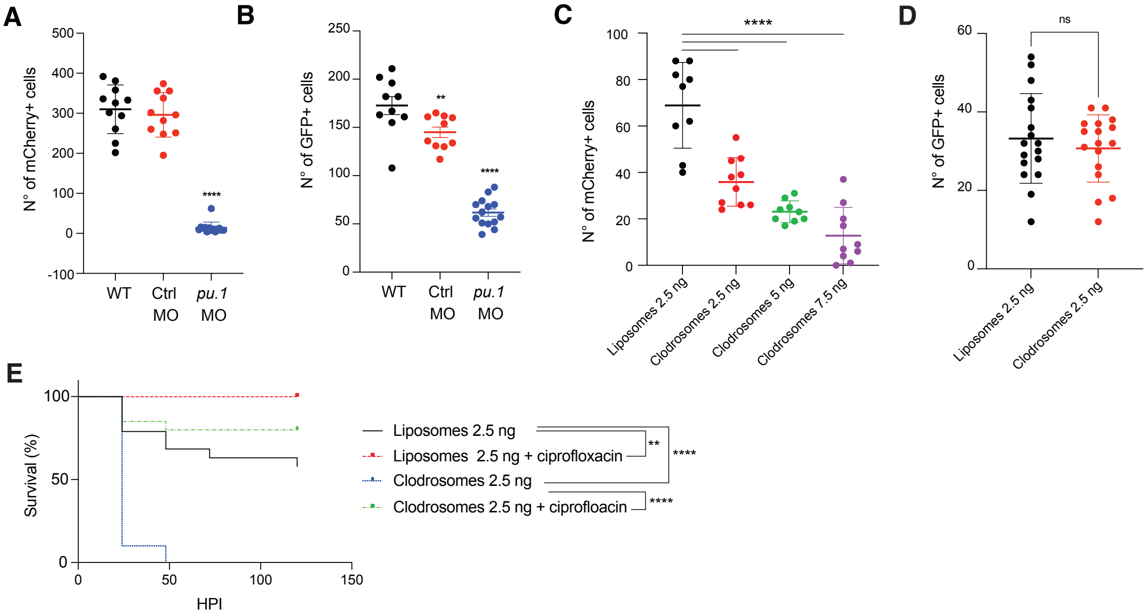


**fig. S3. Depletion of innate immune cells in zebrafish larvae.** Macrophage ((**A**) *mpeg1:mcherry*) and neutrophil ((**B**) *mpx:gfp*) transgenic reporter embryos were injected with either *pu.1* MO or control MO at the single cell stage. *pu.1* morphants showed a significant decrease in whole body macrophage (**A**) and neutrophil (**B**) cell numbers by 3 dpf measured by epifluorescence imaging microscopy. Data shown is the mean +/- SD with N≥ 10 larvae per group and statistical significance determined by One-way ANOVA followed by Dunnett's multiple comparison test with adjusted P values: **, P=0.0098; ****, P<0.0001. (**C**-**D**) Clodrosomes selectively deplete macrophages. Double transgenic *mpeg1:mCherry/mpx:gfp* embryos were injected in the duct of cuvier at 2 dpf with control liposomes or liposomes containing clodronate (clodrosomes) and at 3 dpf (**C**) mCherry+ (N=9-10 embryos/ condition; ****, P<0.0001; 1-way ANOVA with Dunnet's multiple comparisons test) or (**D**) GFP+ cells (N=17 embryos/ condition; two-tailed unpaired t-test) were quantified in the embryos' tail region by epifluorescence microscopy. (**E**) 3 dpf larvae were infected with *S.* Typhimurium (445 +/- 25CFU), treated with ciprofloxacin and monitored over time for survival. Clodrosome mediated depletion of macrophages has no effect on ciprofloxacin treatment (N=18-20 embryos/ condition; **, P<0.0012; ****,P<0.0001; log rank test). Data are representative of 3 independent experiments. dpf; days post fertilization; MO, morpholino; HPI, hours post infection.

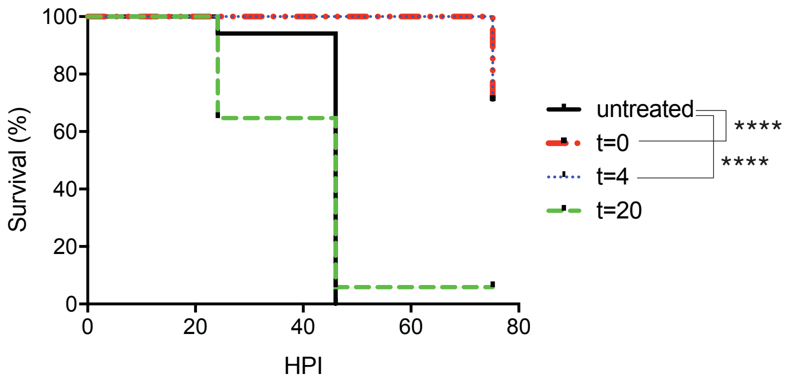


**fig. S4. 3-HK loses full efficacy when administered at 20 hpi.** Larvae (72 hpf) were infected with 346 +/- 166 CFUs of *S.* Typhimurium and treated with 50 uM 3-HK at indicated time points post infection and monitored over time for survival (N=17 embryos/ condition; ****, P<0.0001; log rank test). hpf, hours post fertilization; hpi, hours post infection.

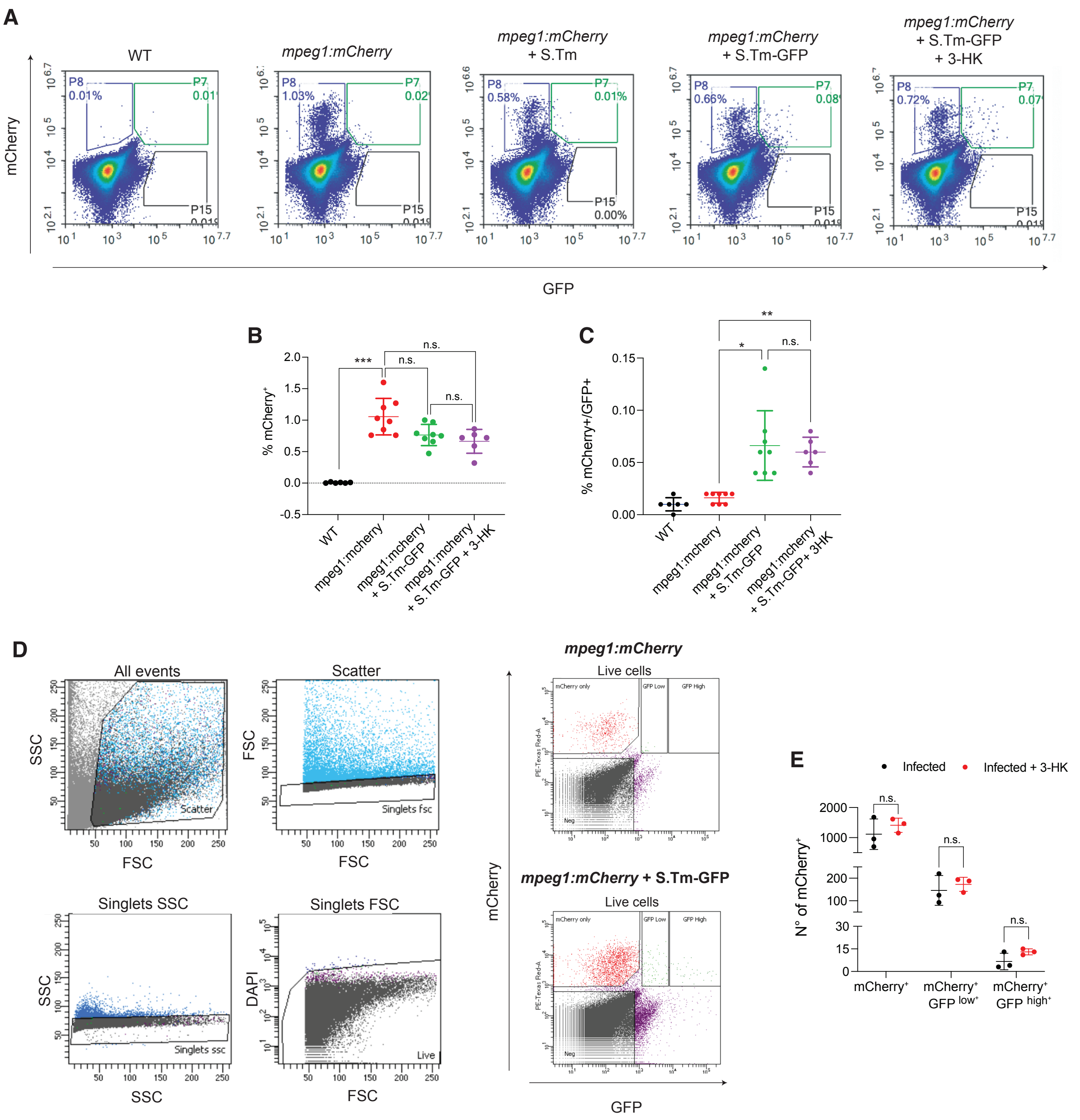


**fig. S5. 3-HK treatment does not lead to differences in the number of viable macrophages or macrophages containing bacteria at 4 hpi.** (**A**) Gating strategy utilized to identify the macrophage population invaded with bacteria in whole embryo single cell suspensions. *mpeg1:mCherry* larvae were infected with *S.*Typhimurium-GFP and at 4 hpi single cell suspensions from groups of 5 animals were prepared and analyzed by flow cytometry. The different conditions indicated above the flow plots were utilized to identify *S.*Typhimurium-GFP (S.Tm-GFP) invaded macrophages. (**B**) Quantification of mean +/- SD of viable mCherry$^+$ cells from S.Tm-GFP (665 +/- 40CFU) infected fish with and without 50 uM 3-HK treatment (6-8 biological replicates per condition). Infection or 3-HK treatment does not lead to changes in the number of viable macrophages measured at 4 hpi (mixed model with Tukey's multiple comparisons test. ***;P=0.0003, N = 3 independent experiments). (**C**) Quantification of mean +/- SD viable mCherry$^+$/GFP$^+$ double positive cells in wild type larvae (WT), macrophage reporters, macrophage reporters infected with S.Tm-GFP with and without 50 uM 3-HK treatment (6-8 biological replicates per condition). Infection or 3-HK treatment does not lead to changes in the number of macrophages associated with bacteria measured at 4 hpi (mixed model with Tukey's multiple comparisons test. **, P=0.0015; *, P= 0.0129. N = 3 independent experiments). (**D**) Gating strategy utilized to FAC sort populations of macrophages (mCherry$^+$) or macrophages asociated with bacteria (mCherry$^+$/GFP$^{low+}$ and mCherry$^+$/GFP$^{high+}$) at 4 hpi. *mpeg1:mCherry* larvae were infected with S.Tm-GFP (662 +/- 13 CFU), treated or untreated with 50 uM 3-HK, and at 4 hpi mCherry$^+$ cells were sorted based upon GFP emission as indicated, (**E**) and mean +/- SD viable mCherry$^+$ cells were quantified for each category (3 biological replicates per condition, two-way ANOVA with Šidák multiple comparison's test. N = 3 independent experiments).

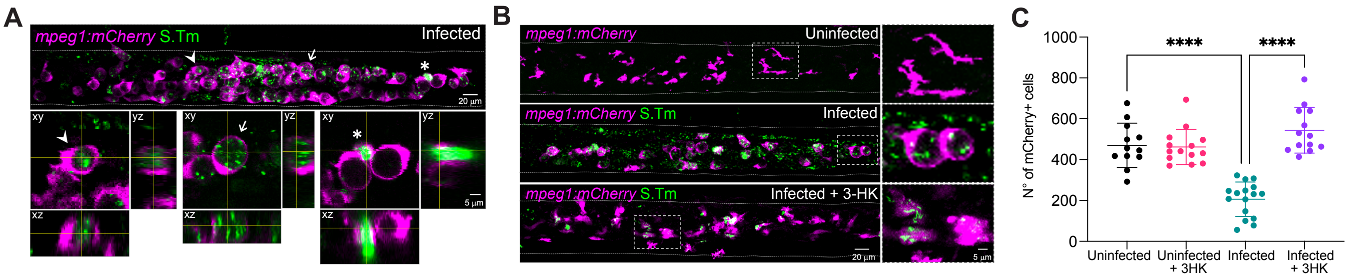


**fig. S6. Multiple *S.* Typhimurium cells can be identified inside macrophage vacuoles in infected zebrafish embryos.** (**A**) Pseudocolored representative maximum intensity projection confocal z-stack image of a *S*. Typhimurium (S.Tm-GFP) infected *mpeg1:mCherry* fish caudal vein at 20 hpi. Enlarged images from the top panel show single confocal sections with z projections for selected cells displaying macrophages with S.Tm-GFP inside vacuoles (*i.e,* arrow and arrow head) and macrophages with reduced volume and extracellular release of S.Tm (*i.e.,* asterix). (**B**) Representative maximum intesity projection of z-stack confocal images of the caudal vein of uninfected, S.Tm-GFP infected, and S.Tm-GFP infected and 50 uM 3-HK treated *mpeg1:mCherry* larvae at 20 hpi. Macrophages from 3-HK treated animals show smaller *S.*Typhimurium containing vacuoles. Right panels show enlarged images of individual cells from demarked sections. (**C**) *S.* Typhimurium infected (347+/- 27 CFU) or uninfected larvae were treated or untreated with 50 uM 3-HK and live imaged at 20 hpi to enumerate mCherry+ cells through whole fish. *S.*Typhimurium infection leads to a reduction in macrophage cell numbers and 3-HK treatment reverses this effect (one-way ANOVA, ****, P<0.0001; **, P=0.0079, Tukey's multiple comparisons test). hpi; hours post infection.

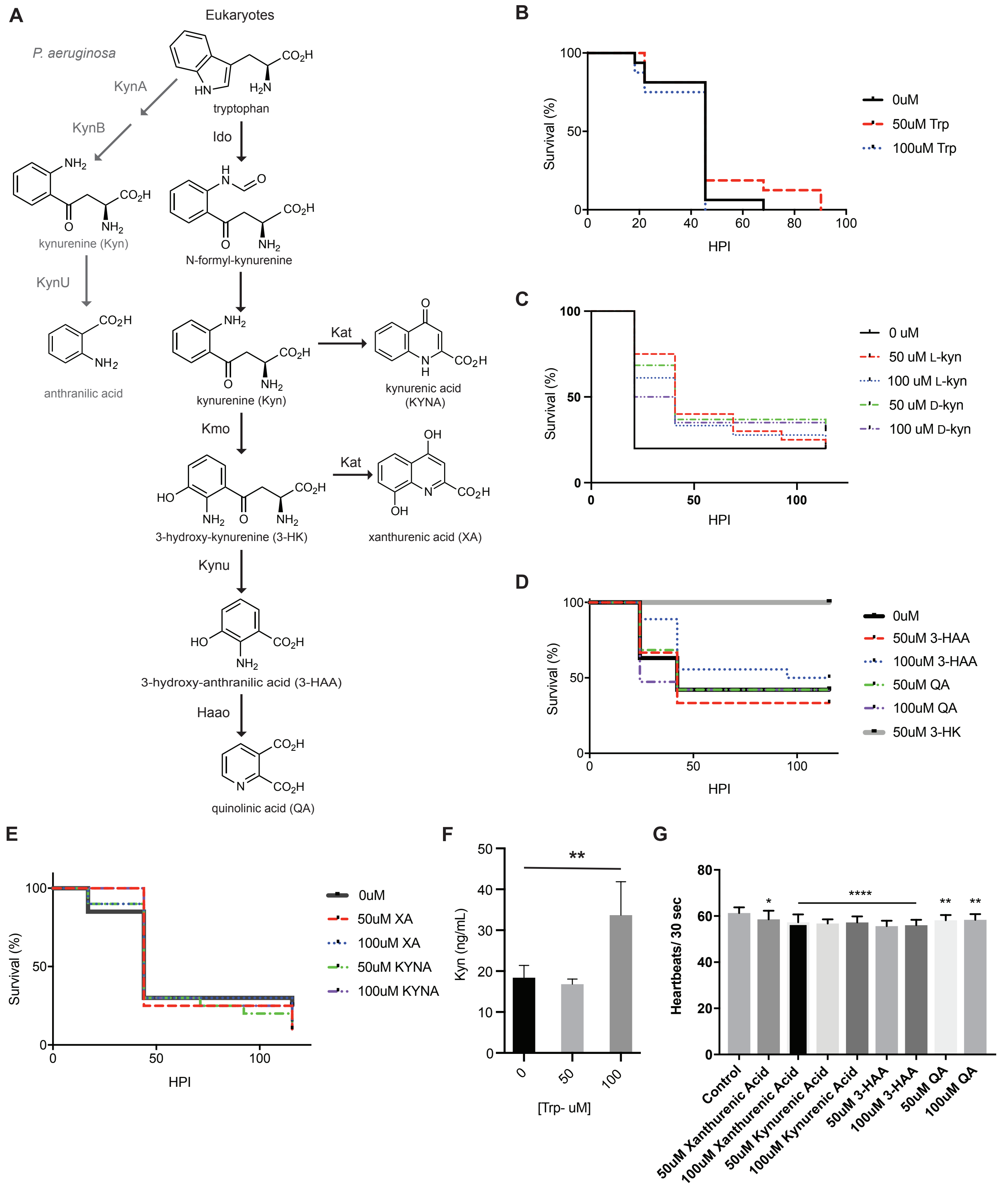


**fig. S7. 3-HK is the only KP metabolite that promotes survival following infection.** (**A**) The Kynurenine pathway in *P. aeruginosa* and in eukaryotes. (**B**-**E**) Larvae (3 dpf) were inoculated with *S.* Typhimurium ((**B**) 453 +/- 106 CFU; (**C**) 350 +/- 42 CFU; (**D**) 229 +/- 81 CFU; and (**E**) 313 +/- 80 CFU) and immersed in indicated concentrations of (**B**) tryptophan; (**C**) L- or D-kynurenine (L- or D-Kyn); (**D**) 3-hydroxy-anthranilic acid (3-HAA), quinolinic acid (QA), or 3-HK; or (**E**) xanthurenic acid (XA) or kynurenic acid (KYNA) and monitored over time for survival. Only 3-HK application had a significant effect on larval survival following infection ((**D**), P=0.0034, Log-rank test; N=16-20 larvae/ group). (**F**) Tryptophan penetrates fish tissues to effect kynurenine levels. Larvae (3 dpf) bathed in tryptophan overnight show a statistically significant increase in kynurenine production by ELISA. Data shown is the mean +/- SD of two combined biological replicates (N=135 larvae/ group). (**G**) 3-hydroxy-anthranilic acid (3-HAA), quinolinic acid (QA), kynurenic acid (KYNA) and xanthurenic acid (XA) penetrate fish tissues to effect heartrate when larvae (3 dpf) were immersed overnight in indicated concentrations. Data shown is the mean +/- SD of two combined biological replicates (N= 13 larvae total/ group except the control group where N=26). Statistical significance was determined by One-way ANOVA ((**F**), P=0.0019; (**G**), P<0.0001) followed by Dunnett's multiple comparison test with adjusted P values: (**F**), P= 0.0039; (**G**) *, P=0.0166; ****, P=0.0001; **, P=0.0043 (50uM QA) and P= 0.0099 (100uM QA).

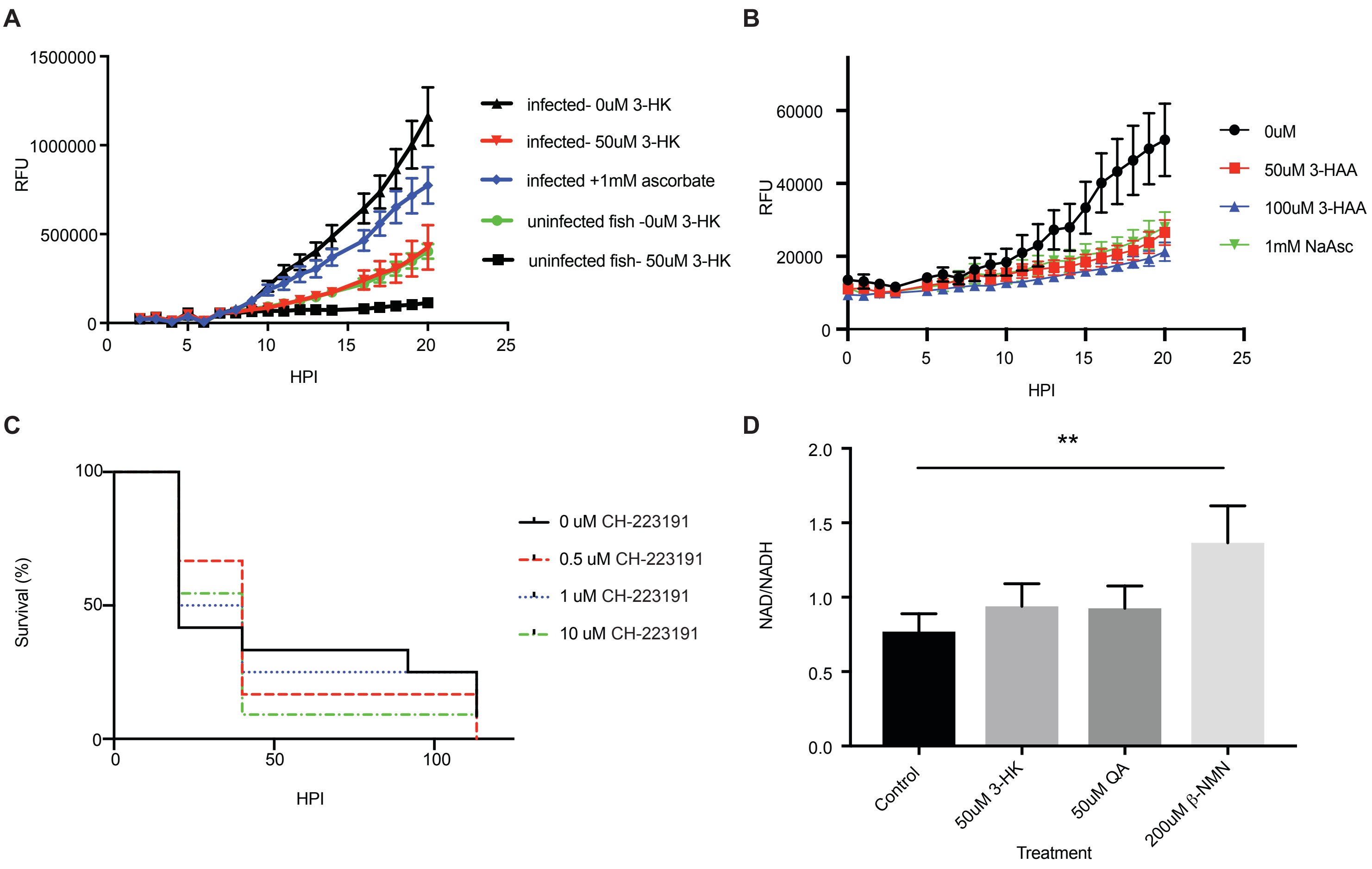


**fig. S8. 3-HK application decreases ROS accumulation, AHR antagonism does not affect survival and 3-HK has no effect on NAD+ levels.** (**A, B**) 3-HK (**A**) or 3-HAA (**B**) immersion results in less ROS accumulation over time in both SL1344-infected (**A**, **B**) and uninfected larvae (**A**). ROS accumulation was determined by bathing larvae in the ROS indicator dye CM-H2DCFDA and monitoring fluorescence over time. Each point indicates the mean and SEM of 6-12 larvae/ condition (1 mM ascorbate was used as a positive control for antioxidant activity). (**C**) Small molecule antagonism of the aryl hydrocarbon receptor (AHR) does not significantly impact survival following infection. Larvae (3 dpf) were inoculated with *S.* Typhimurium (413 +/- 64 CFU) and immersed in indicated concentrations of the AHR antagonist CH-223191 and monitored over time for survival. (**D**) 3-HK does not increase NAD+ levels in fish. Larvae (3 dpf) were immersed in 3-HK or the downstream metabolite QA overnight compared to the positive control, β-NMN, which is an NAD precursor that boosts NAD+ levels through the salvage pathway and NAD/NADH ratios were measured using the NAD/NADH-Glo Assay. The data are the mean +/- SD of N=3 larvae/ condition. Statistical significance was assessed by one-way ANOVA ($p=0.0161$) followed by Dunnett's multiple comparison test (adjusted $P= 0.0080$). The data are representative of 3 independent experiments.

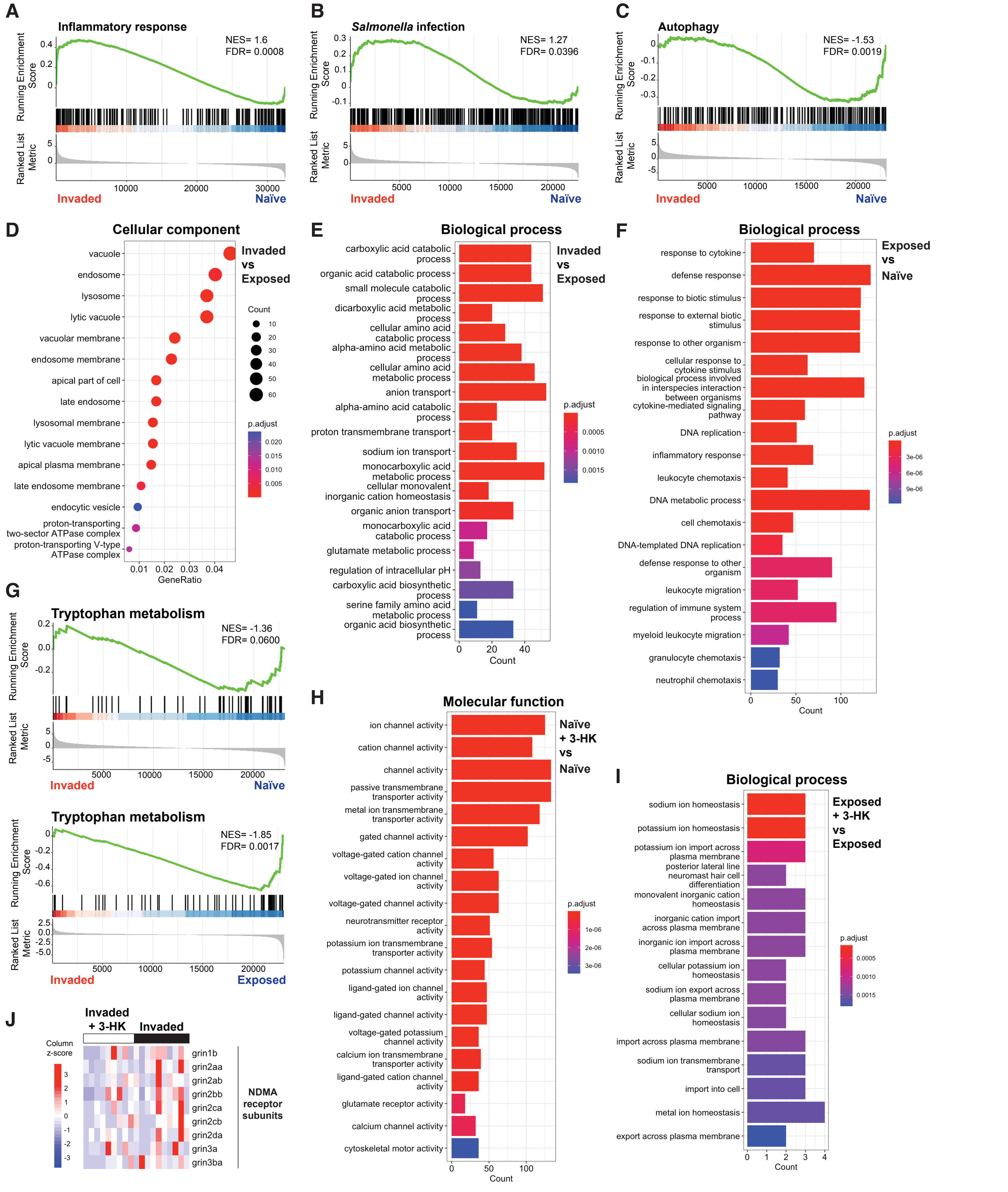


**fig. S9. Significantly enriched gene ontology biological processes identified by transcriptomic analyses of macrophages isolated from *S.* Typhimurium infected embryos treated or untreated with 3-HK and measured at 4 hours post infection.** (**A**-**C**) Gene Set Enrichment Analysis (GSEA) plots obtained from functional analysis of differentially expressed genes from *S.* Typhimurium invaded versus naïve macrophages show activation of gene sets associated with (**A**) inflammatory response (**B**) *Salmonella* infection and suppresion of gene sets associated with (**C**) autophagy. (**D**-**E**) Gene ontology plots showing enrichment of celluar components (**D**) and biological processes (**E**) for the genes differentially expressed between *S.* Typhimurium invaded versus exposed macrophages. (**F**) Gene ontology plot showing showing enrichment of biological processes for the genes differentially expressed between *S.* Typhimurium exposed versus naïve macrophages. (**G**) GSEA plots showing supression of tryptophan metabolism in *S.* Typhimurium invaded versus naïve macrophages (upper panel) or *S.* Typhimurium invaded versus exposed macrophages (lower panel). (**H**) Gene ontology plot showing enrichment of molecular functions for the genes differentially expressed between *S.* Typhimurium exposed versus naïve macrophages. Statistical significance (p.adj) for the enriched terms is displayed and color-coded. (**I**) Heatmaps showing NMDA receptor codifying genes differential expression between *S.* Typhimurium invaded macrophages isolated from 3-HK treated animals versus *S.* Typhimurium invaded macrophages isolated from vehicle treated animals. Statistical analysis shows no significant differential expression with a significance cut-off of p.adj < 0.05 for any gene. NES: normalized enrichment score; FDR: false discovery rate.

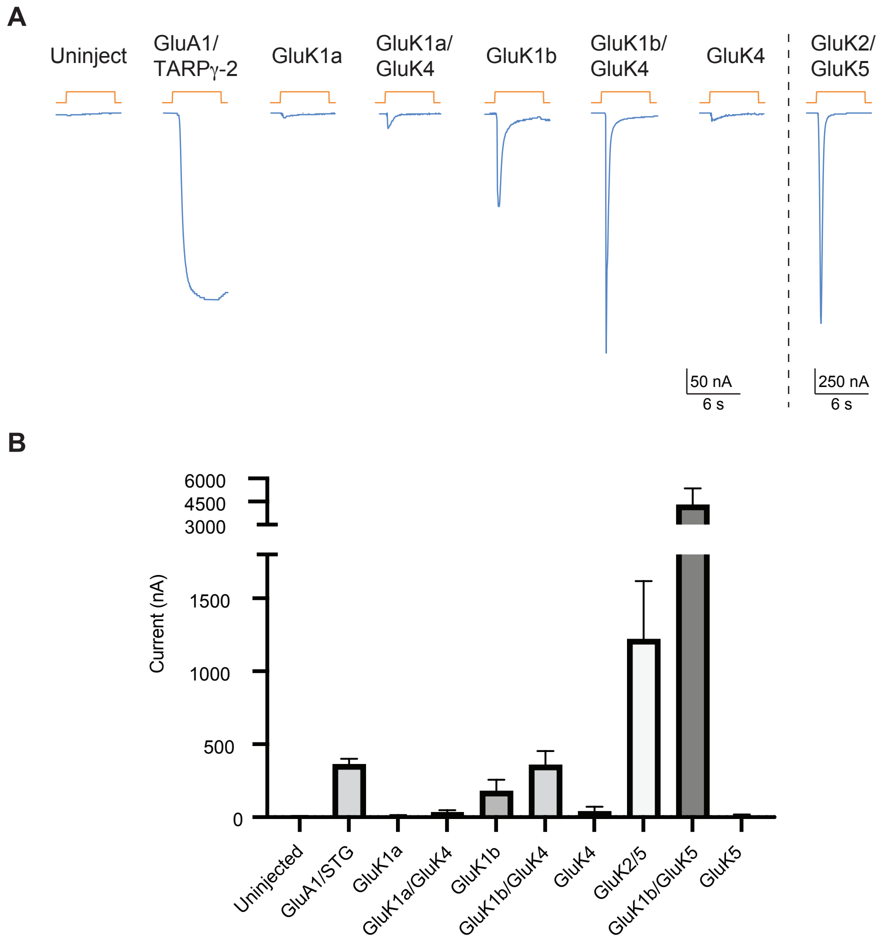


**fig. S10. The *D. rerio* KAR subunit GluK1b forms glutamate-gated channels in heterologous cells alone or in complex with the high affinity subunits GluK4 or GluK5.** (**A**, **B**) Representative traces of glutamate- evoked currents (100 uM; **A**) were observed in oocytes co-expressing *D. rerio* GluK1b (*grik1b*), GluK1b/GluK4 (*grik1b*/*grik4*) or GluK1b/GluK5 (*grik1b*/*grik5*) with murine Neto2 as well as oocytes expressing the murine AMPAR GluA1 with TARPγ-2 or the murine KAR GluK2/GluK5 (*grik2*/ *grik5)* with Neto2. No current was observed in oocytes expressing GluK1a *(grik1a)* alone or with GluK4 *(grik4).* As expected, no obvious currents were observed in oocytes expressing only the high affinity subunits GluK4 *(grik4) or* GluK5 *(grik5)* which require heterotetramerization with GluK1-GluK3. KAR and AMPAR protein subunit names are displayed with the corresponding gene names indicated parenthetically. (**B**) Mean +/- SEM of replicate current measurements (N=5-7).

**Table S1. Summary of Cell Culture Experiments with 3-HK**

| Cells/ lines | J774 | RAW264 | U937 | Murine BMDMs |
|---|---|---|---|---|
| Host Cell Survival | n.s.[1,2,3] | n.s.[2,4] | n.s.[1,2,3] | n.s.[1,4] |
| Bacterial Burden | n.s.[1,2] | n.s.[2] | n.s.[1,2] | |

[1] infections performed with *S.* Typhimurium grown to log phase
[2] infections performed with *S.* Typhimurium grown to stationary phase
[3] assayed with Cell Titer Glo
[4] assayed with LDH assay
n.s.: no significant difference with 3-HK treatment
